# Stimulus selection influences prediction of individual phenotypes in naturalistic conditions

**DOI:** 10.1101/2023.12.07.570273

**Authors:** Xuan Li, Simon B. Eickhoff, Susanne Weis

## Abstract

Understanding individual differences and brain-behaviour relationships is an essential goal of human neuroscience. Recent studies have shown the great potential of naturalistic stimuli, e.g., movie clips, in advancing this pursuit. While the use of naturalistic stimuli attracts increasing interest, the influence of stimulus selection remains largely unclear. In this study, we show that brain activity is generally sensitive to the choice of movie stimuli at both group and individual subject levels. Using sex classification as an example, we demonstrate that brain activity elicited by different stimuli can lead to distinct prediction performance and unique predictive features. The stimuli that yield better classification performance often elicit stronger synchrony of brain activity across all subjects and are mostly derived from Hollywood films with rich social content and cohesive narratives. Our results highlight the importance of stimulus selection and provide practical guidance for choosing appropriate stimuli, opening up new avenues for future studies on individual differences and brain-behaviour relationships.

## Introduction

Understanding interindividual variability in brain function in relation to behaviour is a central goal of human neuroscience. Naturalistic stimuli, e.g., movies or spoken stories, have recently emerged as a promising tool for advancing this goal. They introduce more synchrony in brain activity across subjects relative to unconstrained resting-state conditions (Hasson et al., 2004), mitigating unsystematic noises and improving interpretability of measured brain signals (Vanderwal et al., 2015). They also provide more ecological validity than conventional tasks by engaging the brain in conditions more similar to real-world situations (Sonkusare et al., 2019), offering novel insights into complex cognitive processes, such as hierarchical organisation of memory (Hasson et al., 2015) and naturalistic emotions (Nummenmaa et al., 2012). While most studies have used naturalistic stimuli for investigating brain activity patterns shared across subjects, an increasing number of studies have suggested their utility for studying individual differences. For example, previous studies have found that naturalistic stimuli evoke reliable individual differences in brain activity, promoting individual identification and showing associations with stable personal traits and mental disorders (Byrge et al., 2015; Campbell et al., 2015; Finn et al., 2020; Vanderwal et al., 2017).

Simultaneously, machine learning techniques have become an increasingly important tool in exploring individual differences and brain-behaviour relationships. They allow researchers to exploit nuanced individual differences in brain activity patterns to make individual-level predictions on novel samples, and have successfully been applied to predict individuals’ cognitive ability, personality and clinical symptoms (Cai et al., 2020; Dubois et al., 2018; Gabrieli et al., 2015; Sripada et al., 2020). Machine learning approaches not only facilitate our understanding of neural correlates of human cognition and behaviour (Rosenberg et al., 2018) but may also advance clinical research and applications, e.g., biomarker discovery for mental disorders and personalised treatments (Bzdok et al., 2020; Woo et al., 2017). While most studies have applied machine learning for phenotype prediction on brain activity signals during rest or conventional tasks, recent studies have shown that naturalistic conditions outperform resting or task conditions in prediction performance (Finn & Bandettini, 2021; Li et al., 2023), with significant potential for clinical applications (Eickhoff et al., 2020). Together, conducting phenotype prediction on brain data in naturalistic conditions is emerging as a valuable tool for understanding individual differences and brain-behaviour relationships.

Despite the promise, one often underestimated concern in naturalistic studies is the impact of stimulus selection. Choosing an appropriate stimulus is a fundamental consideration in experimental design, yet most naturalistic studies and datasets so far have not provided sufficient justifications for selecting specific stimuli. One main reason for this is the current lack of a systematic approach for stimulus selection. Furthermore, it is expected that brain activity would vary across different stimuli, as the brain is known to respond differentially to various inputs (Hasson et al., 2010). However, how stimulus selection influences investigations of individual differences and phenotype prediction remains largely unclear. Some studies have suggested that stable individual traits play a more dominant role than external stimuli in modulating brain activity, resulting in broadly similar representations of individual differences and phenotype prediction performance across different stimuli (Vanderwal et al., 2017; Gao et al., 2020; Tian et al., 2021). Conversely, other studies have observed distinct representations of individual differences and phenotype prediction performance across different stimuli, suggesting a significant impact of stimulus selection (Finn & Bandettini, 2021; Kröll et al., 2023; Li et al., 2023). Therefore, gaining a better understanding of the influence of stimulus selection will benefit naturalistic studies in both the experimental design for data acquisition and result interpretation.

In this present study, we investigated the influence of stimulus selection from three different aspects (Fig. 1A). Specifically, we examined 1) variations in brain states elicited by different stimuli, 2) disparities in phenotype prediction performance across stimuli, and 3) features of the stimuli that could potentially influence brain states and their utility for phenotype prediction. To investigate the influence of stimulus selection on brain states, we focused on stimulus-evoked brain activity, i.e., brain activity that is largely time-locked to the stimulus and consistent across subjects, thus often characterised based on inter-subject synchrony (Hasson et al., 2004; Nastase et al., 2019). Compared to spontaneous brain activity, another component of brain signals in naturalistic conditions, stimulus-evoked brain activity is often of more interest to researchers and has been shown to be closely linked to behaviour (Finn et al., 2020). For the same reason, we used our recently proposed approach, the topography-based predictive framework (TOPF), for phenotype prediction (Li et al., 2023). This approach allows us to capture individual differences in stimulus-evoked brain activity in a data-driven way and uses them as features for prediction of individual phenotypes in a machine learning framework (Fig. 1B).

**Fig. 1:**
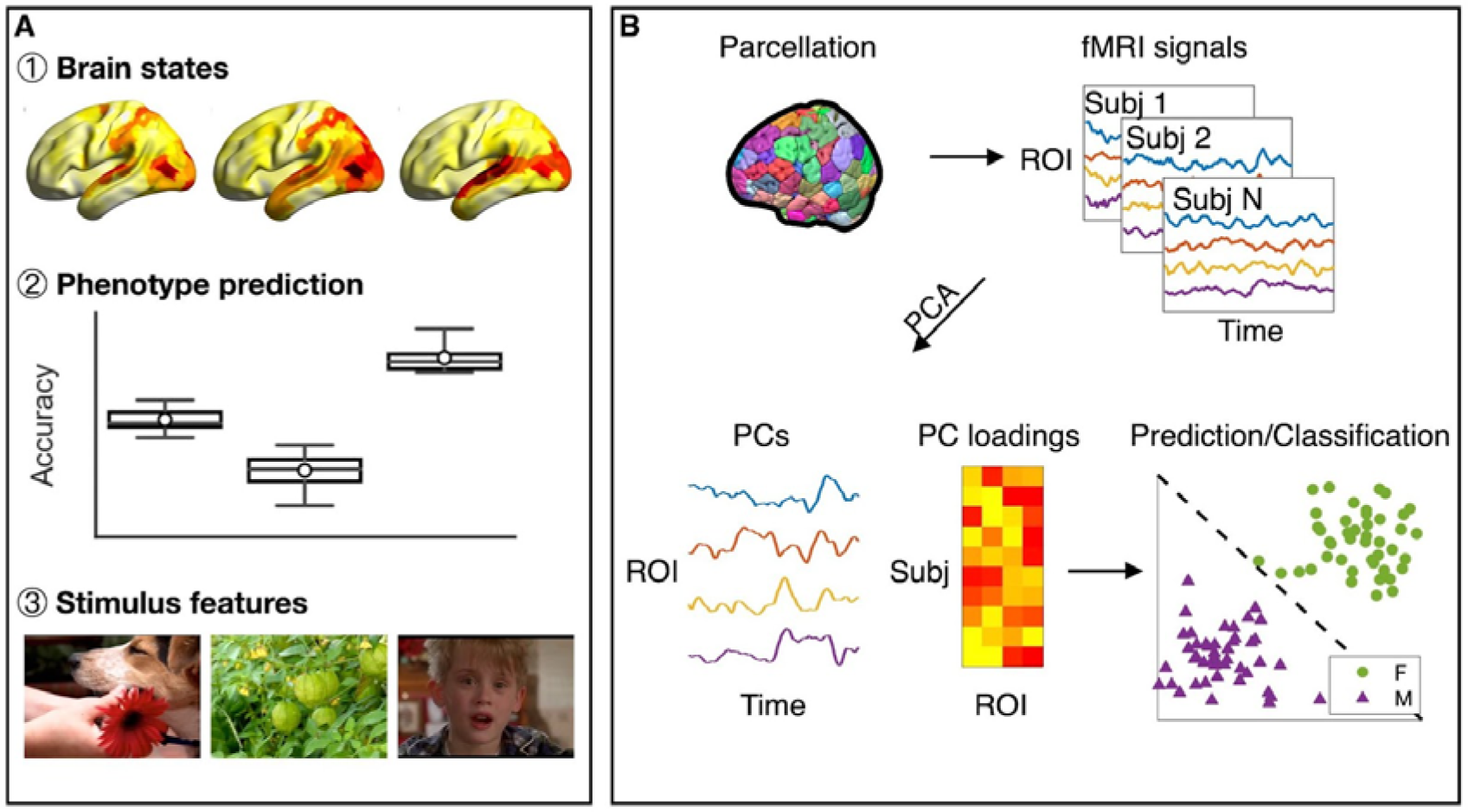
Overview of research questions and the pipeline for phenotype prediction using TOPF. **A**) In this study, we investigate the influence of stimulus selection on studies of individual differences and phenotype prediction during naturalistic conditions from three aspects. Specifically, we ask Ill whether different movie stimuli elicit different brain states; Ill whether phenotype prediction performance varies across different movie stimuli; Ill which features of movie stimuli could potentially influence their utility for studying individual differences and phenotype prediction. **B**) Schematic illustration of the TOPF method. Image reproduced from (Li et al., 2023), copyright ©2023 Xuan Li. A principal component analysis (PCA) is applied to fMRI time series across subjects for each region of interest (ROI) separately. The derived PCs represent shared responses across subjects, with PC loadings reflecting individual expression levels of the shared responses. The PC loadings are used as features for phenotype prediction (see “Methods” for detailed descriptions of the method).

Among various types of naturalistic stimuli, we focused on movie stimuli, a popular choice of stimuli due to their complexity, multimodality and capability of eliciting brain responses broadly across the whole brain (Hasson et al., 2010; Hasson, Yang, et al., 2008). We used the human connectome project (HCP) dataset (Van Essen et al., 2013), which provides functional magnetic resonance imaging (fMRI) data acquired during watching a variety of movie clips and a relatively large number of samples for machine learning analysis. For phenotype prediction, we limited our analyses to a single phenotype, biological (i.e., birth-assigned) sex, due to its robust nature and well-demonstrated associations with various higher-order cognitive functions (Miller & Halpern, 2014; Weis et al., 2020). We examined whether different movie stimuli can elicit distinct group-level brain states, measured by inter-subject synchrony, and different representations of individual differences in brain activity. We next tested whether performance for phenotype prediction varies across different stimuli, and in particular how this is related to inter-subject synchrony. Finally, we delved into features of the movie clips themselves to explore potential origins of differences across these stimuli in the evoked brain activity and utility for prediction.

## Results

We used fMRI data from all available subjects who completed watching all movie clips (HCP S1200 release), yielding a sample of 178 subjects. We considered movie clips that were presented only once in the HCP study and had a length of over 2 mins, resulting in 13 movie clips covering various topics (2 - 4 mins; Supplementary Table1). All movie clips were truncated to the length of the shortest movie clip (132 TRs, i.e., 2:12) except in “Control analyses” where the impact of different clip lengths was assessed.

### Different movie clips elicit distinct levels of inter-subject synchrony

Measuring shared brain response across subjects is an important approach to examine stimulus-evoked brain activity in naturalistic studies (Hasson et al., 2004; Nastase et al., 2019; Simony et al., 2016). Such shared response is often characterised by inter-subject synchrony. The core idea of such an approach is that any consistent, synchronous fluctuations of brain activity across subjects is a result of processing of the same stimulus. In practice, the level of inter-subject synchrony is also often used to reflect how much control a stimulus exerts over subjects‘ brains (Hasson, Landesman, et al., 2008). To test whether the brain state elicited during watching different movie clips varies at a group level, we computed the inter-subject synchrony levels of brain activity for each movie clip.

Specifically, we extracted mean time series over voxels within each of 400 cortical (Schaefer et al., 2018) and 36 subcortical (Fan et al., 2016) regions of interest (ROIs). For each ROI, we performed a principal component analysis (PCA) to the fMRI time series across subjects and represented the shared response across subjects by the first principal component (PC1) of these time series (Di & Biswal, 2022; Li et al., 2023). The inter-subject synchrony was quantified as the variance explained by PC1 for each ROI. A larger amount of explained variance indicates that temporal fluctuation patterns of brain activity are more consistent across subjects, i.e., a higher level of inter-subject synchrony.

Fig. 2A shows the inter-subject synchrony values of all ROIs for each movie clip. A one-way ANOVA revealed significant differences between movie clips (p<0.0001). “Inception” and “dreary” achieved the highest (PC1 variance: 20.9±12.8%) and lowest (PC1 variance: 7.9±4.0%) average inter-subject synchrony level, respectively. Fig. 2B shows a brain map of inter-subject synchrony averaged over all movie clips (see Supplementary Fig. 1 for maps of individual movie clips). Consistent with previous studies (Hasson et al., 2004), the temporal, occipital and parietal cortices exhibited remarkably stronger inter-subject synchrony than the other brain regions.

**Fig. 2:**
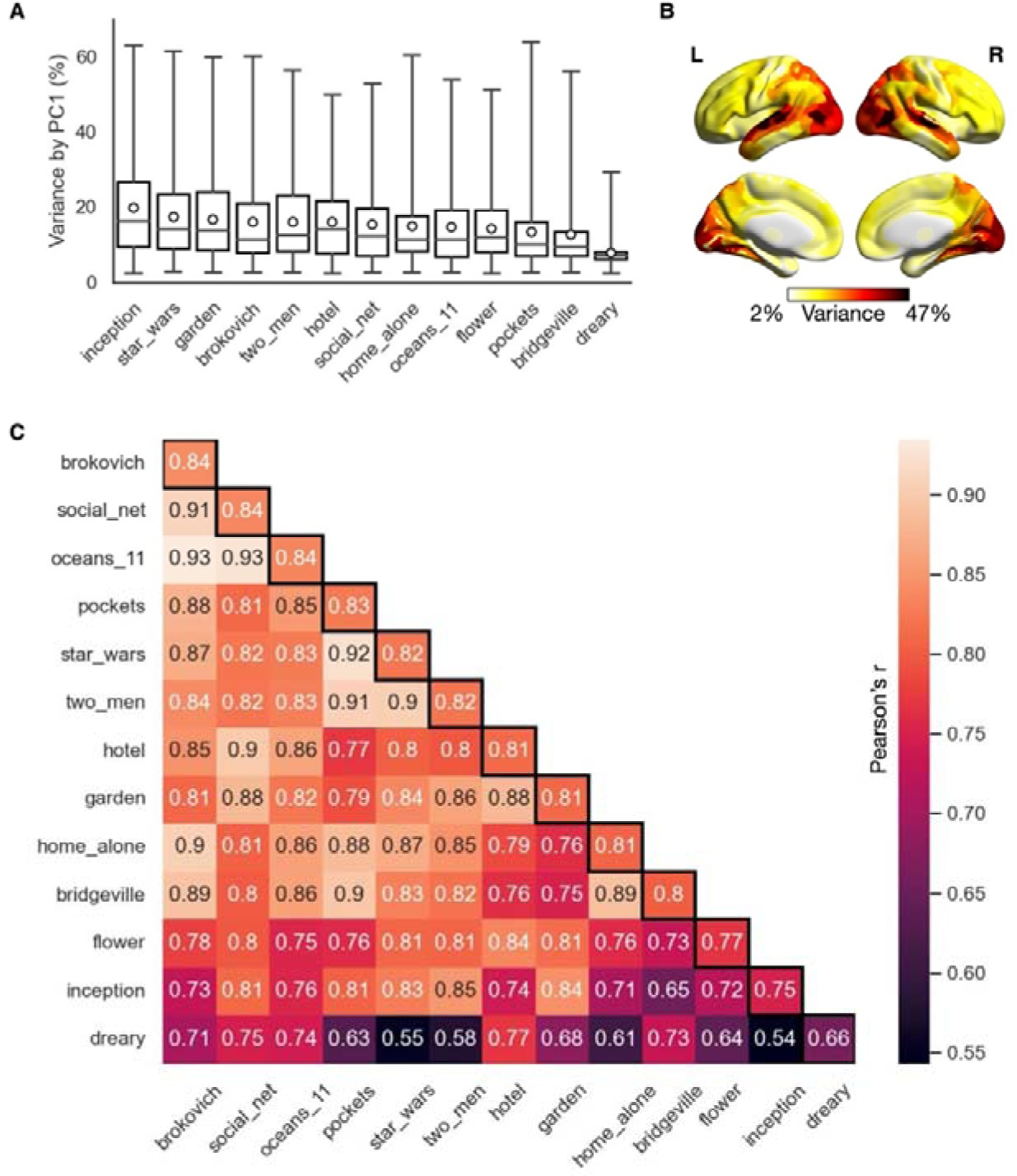
Different movie clips can elicit different levels and spatial patterns of inter-subject synchrony. **A**) Inter-subject synchrony across all ROIs for individual movie clips, measured as the variance explained by the PC1 of the fMRI time series across subjects. The three bars of each box from top to bottom indicate the third quartile, median and first quartile, respectively. The circle in the middle represents the mean. The upper and lower whiskers represent the maximum and minimum, respectively. Movie clips are ordered according to the mean inter-subject synchrony value over ROIs. **B**) Spatial pattern of inter-subject synchrony across the whole brain. For each ROI, the inter-subject synchrony value was averaged over all movie clips. The light-dark colour bar indicates the inter-subject synchrony value from low to high. **C**) Similarity (Pearson’s r) between movie clips in spatial patterns of inter-subject synchrony. Movie clips are ordered according to their average similarity to the other clips (diagonals in black boxes). High and low similarity values are indicated by light and dark colours, respectively.

To assess the similarity between movie clips in the spatial patterns of inter-subject synchrony, we correlated the inter-subject synchrony values of all ROIs between each pair of clips, resulting in a 13 by 13 correlation matrix (Fig. 2C). In general, the spatial pattern of inter-subject synchrony was consistent across the clips, with the mean similarity over all clip pairs achieving r = 0.80±0.05. Notably, “dreary” elicited a highly different pattern from the other movie clips (mean r = 0.66±0.08). We found that while the other movie clips evoked strong inter-subject synchrony extensively across the sensory cortices, “dreary” primarily evoked inter-subject synchrony within the visual cortex (although the values were also much lower than those of the other clips). This unique pattern of “dreary” might be due to the fact that “dreary” is the only movie clip that contains only images of natural scenery without any human images and speech. Overall, we found that the brain state varied largely across different movie stimuli in terms of both the degree and the spatial pattern of inter-subject synchrony.

### Movie stimuli modulate representations of individual differences in evoked brain activity

We next investigated how different movie clips influence brain activity at an individual subject level. Individual differences in response to the same stimulus in brain activity are often represented by unique expression levels of the shared response (Di & Biswal, 2022; Finn et al., 2020). Specifically, for each ROI we operationalised individual expression levels of shared responses as subject-wise loadings onto the PC1 of the fMRI time series across subjects (Li et al., 2013). A high loading value indicates that brain activity of a given subject closely resembles the shared response. To assess how individual expression levels change across clips, we correlated the PC1 loadings of all subjects between each pair of movie clips within each ROI, resulting in a 13 by 13 correlation matrix for each ROI. A high correlation value indicates that the representation of individual differences is highly similar between movie clips.

Fig. 3A shows the correlation averaged over all clip pairs for each ROI. Overall, the average similarity across clips varied largely across the whole brain (r = 0 - 0.47), with sensory cortices achieving the highest values and motor cortex and subcortical regions achieving the lowest values (Gao et al., 2020; Li et al., 2023). Fig. 3B shows the correlations between all pairs of movie clips of all ROIs, grouped by network (Yeo et al., 2011). Similarly, representations of individual differences were more similar across movie clips in visual, temporal-parietal and dorsal attention networks compared to the other networks. These results suggest that in certain brain systems the representation of individual differences was relatively stable across different stimuli and insensitive to changes in the specific content of stimuli.

**Fig. 3:**
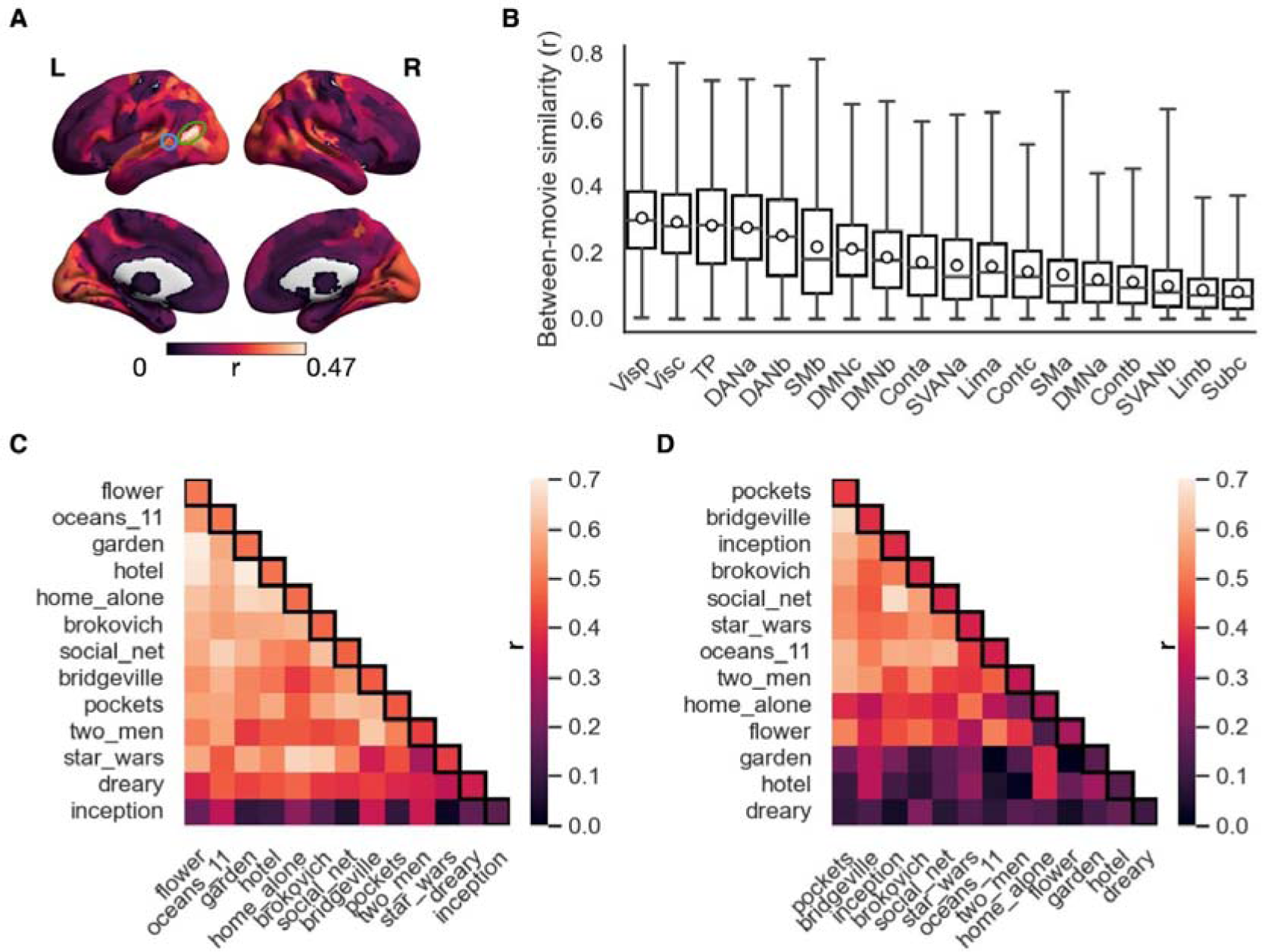
Representations of individual differences in evoked brain activity vary across different movie clips. **A**) Spatial pattern across the whole brain of the similarity in the representation of individual differences across movie clips. Individual differences in evoked brain activity were represented by individual expression levels of shared responses, thereby computed as the subject-wise loadings of the PC1 of fMRI time series across subjects. The value of each ROI represents the average correlation coefficient (Pearson’s r) of PC1 loadings across subjects over all movie clip pairs. High and low values are indicated by light and dark colours, respectively. **B**) The correlations of all ROIs and movie clip pairs grouped by network. Boxplot convention is consistent with that used in Fig. 2A. **C**) Correlation matrix for the ROI (marked in a green circle in A) achieving the highest average between-movie similarity. **D**) Correlation matrix for the ROI (marked in a blue circle in A) achieving the highest standard deviation of the between-movie similarity. Movie clips are ordered according to their average similarity to the other clips (diagonals in black boxes).

In addition, the similarity of the representation of individual differences across clips also highly depended on which movie clips were being compared (r = 0 - 0.79). For example, Fig. 3C shows the between-movie correlation matrix for a ROI in the left temporal parietal junction (TPJ), a key region in the dorsal attentional network (green circle in Fig. 3A). The representation of individual differences was most stable across all movie clips in this ROI (r = 0.47±0.16). We found that “inception” and “dreary” were on average least similar to the other movie clips in this ROI (diagonals in Fig. 3C; inception: r = 0.16±0.11; dreary: r = 0.36±0.13). Notably, these two movie clips were also the ones that elicited the strongest and lowest inter-subject synchrony, respectively (Fig. 2A). These results suggest that the brain might be in certain unique states during watching these two movie clips that resulted in distinct representations of individual differences.

As an additional example, Fig. 3D shows the between-movie correlation matrix of a ROI in the left superior temporal gyrus (STG) (blue circle in Fig. 3A). STG is known for its crucial role in language and speech processing (Mesgarani et al., 2014). This ROI exhibited the largest standard deviation of the similarity across clips (r = 0.38±0.20). In this ROI, we found that “garden” (r = 0.16±0.11), “hotel” (r = 0.16±0.11) and “dreary” (r = 0.11±0.06) were least similar to the other movie clips in the representation of individual differences (Fig. 3D). An interesting common characteristic of these movie clips is that they were all intentionally designed to minimise dialogue content. These results demonstrate that changes in specific attributes of movie stimuli can lead to large variations in the representation of individual differences.

### Utility for sex classification depends on the choice of movie stimuli

Given that brain states varied while watching different movie stimuli, we next investigated whether these differences extended to their utility in phenotype prediction. To do this, we applied the TOPF method (Li et al., 2023) to each movie clip separately to predict subjects’ biological sex (Fig. 1B). For each subject, we used the individual expression levels of shared responses (i.e., PC loadings) from all ROIs as features for sex classification. In our main analysis, we used only PC1 loadings, resulting in 436 features. We tested different choices of feature spaces in “Control analyses”.

We employed a support vector machine (SVM) classifier with a radial basis function (RBF) kernel for sex classification (Weis et al., 2020). Classification performance was evaluated via a 10-fold cross validation (CV). In each CV fold, all subjects were split into a training set and a test set. We fitted the classifier on the training set and applied it to predict sex of subjects in the test set. To prevent data leakage, we ensured that training and test subjects were fully separated before feature extraction and that subjects from the same family were all included in either the training or the test set. We repeated the procedure 10 times and measured classification performance as the mean balanced accuracy over all CV folds and repetitions.

Fig. 4A shows the sex classification performance of all movie clips. The accuracy significantly exceeded the chance level (permutation-based p<0.05; FDR corrected) for all movie clips, except for “dreary”. Classification performance varied largely across movie clips, with “star wars” achieving the highest accuracy (69.8%) and “dreary” the lowest (44.7%) among all clips. Corrected resampled t-tests revealed that the performance of “dreary” was significantly worse than that of most of the other movie clips (p<0.05; FDR corrected; Fig. 4B). Overall, these results suggest that there were considerable differences across the movie clips in the sex classification performance.

**Fig. 4:**
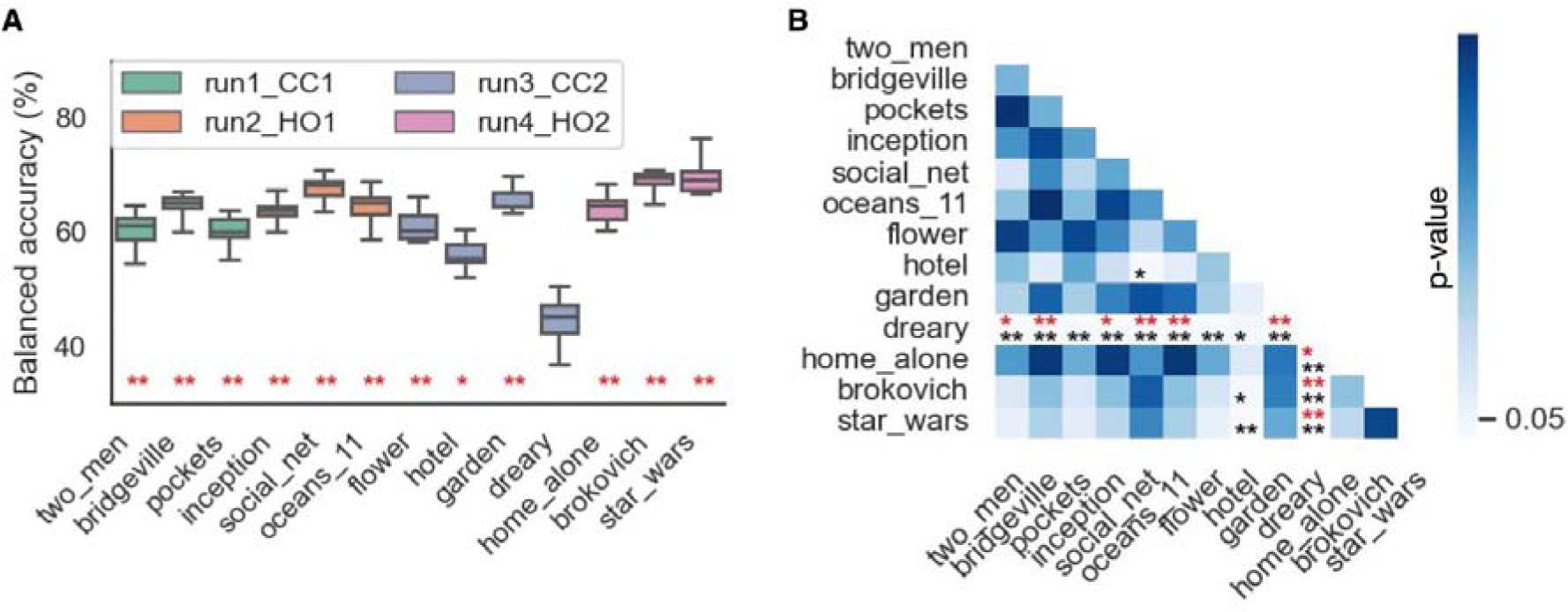
Sex classification performance varies across movie clips. **A**) Sex classification performance (balanced accuracy) of individual movie clips using TOPF. The boxplots show the scores from 10 repetitions. The three bars of each box from top to bottom indicate the third quartile, median and first quartile, respectively. The upper and lower whiskers represent the maximum and minimum, respectively. A permutation test (1000 iterations) was used to evaluate whether the accuracy was significantly above chance. * denotes p<0.05 (FDR corrected). ** denotes p<0.01 (FDR-corrected). CC and HO represent the runs using clips from independent films and Hollywood films, respectively. **B**) Comparison of classification accuracies between movie clips by corrected resampled paired t-tests. * denotes p<0.05. ** denotes p<0.01. P-values before and after FDR correction for multiple comparisons are marked in black and red, respectively.

### Predictive features and models vary across movie clips

Having shown that performance for sex classification varied when using different movie clips, we assessed whether predictive features were different across these movie clips. We computed the permutation importance for each feature separately (Breiman, 2001). Permutation importance of a feature was quantified as the decrease in classification accuracy when the feature was shuffled across subjects. A positive value indicates that the feature is important for sex classification.

Fig. 5A shows the importances (averaged over all movie clips) of all features (i.e., ROIs). Overall, the left prefrontal, temporal and parietal cortices and right STG and dorsolateral prefrontal cortex exhibited high importance for sex classification (Weis et al., 2020). Fig. 5B shows the feature importance maps of two representative movie clips, i.e., “star wars” and “brockovich” (see Supplementary Fig. 2 for maps of all clips). While these two movie clips achieved the highest classification accuracy (“star wars”: 69.8%; “brockovich”: 69.1%; Fig. 4A), we found that proximity in classification performance does not imply similarity in predictive features. The important features spread across the whole brain for “star wars”, while for “brockovich”, they mainly clustered in the dorsolateral prefrontal cortex, parietal and temporal lobes.

**Fig. 5:**
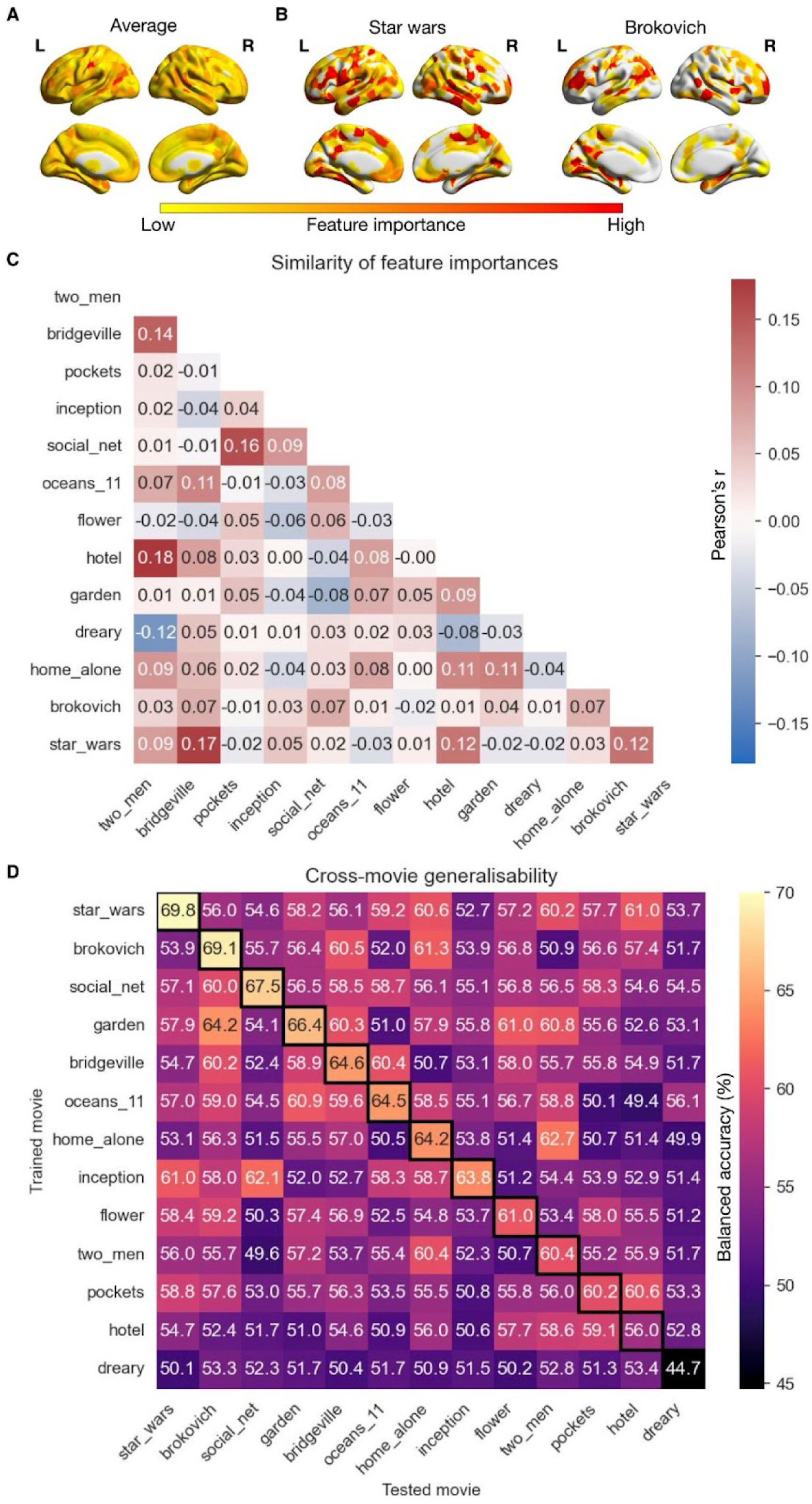
Comparisons of predictive features and models between movie clips. **A**) Feature importance map averaged over all movie clips. **B**) Feature importance maps of two representative movie clips, “star wars” and “brockovich”. For (**A**) and (**B**), permutation importance of each feature was measured as the decrease in balanced accuracy after shuffling the feature across subjects. Negative values were set to zeros for illustration. The yellow-red colour bar represents the importance values from low to high. **C**) Similarity (Pearson’s r) of feature importance maps (Supplementary Fig. 2) for each pair of movie clips. Negative and positive values are indicated by the colours blue and red, respectively. **D**) Generalisability of predictive models across movie clips. Each value in the matrix represents the balanced accuracy obtained when applying the models trained on one movie clip to another clip and averaged over all CV folds. Movie clips are ordered according to their within-movie classification accuracies (diagonals marked in boxes). Lower and higher accuracies are indicated by dark and light colours, respectively.

To quantify the similarity of predictive features between movie clips, we correlated the feature importances of all ROIs between each pair of clips, resulting in a 13 by 13 matrix (Fig. 5C). The similarity was in general low (r = -0.12 - 0.18) across clip pairs, with the similarity between “star wars” and “brockovich” only achieving r = 0.12. These results suggest that brain regions that contributed most to sex classification varied across different movie stimuli.

In a second analysis, we evaluated whether the predictive models can generalise across different movie clips. For each pair of movie clips, we applied the models trained on clip A to predict subjects’ sex from data of clip B. Fig. 5D shows the cross-movie classification performance for all pairs of clips. Movie clips are ordered according to their within-movie classification accuracy (i.e., the diagonals of the matrix). “Star wars” and “dreary” achieved the highest and lowest average cross-movie generalisability, respectively (star wars: 57.7±2.9%; dreary: 51.5±1.2%). We found that the average generalisability to other clips of a movie clip (i.e., row-wise means without diagonals) was highly correlated with its within-movie classification performance (r = 0.89, p <0.0001; Supplementary Fig. 3A). This result suggests that models that achieved higher within-movie classification performance may better capture sex differences related to the general processing of movie stimuli. On the other hand, the generalisability varied largely across movie clip pairs (49.4% - 64.2%), with the values (averaged within each pair of clips over the two directions) significantly correlated with the similarity in feature importance (r = 0.26, p = 0.02; Supplementary Fig. 3B). This suggests that movie clips highlighting similar brain regions for sex classification tended to generalise well to each other. Overall, these results suggest that different movie clips may produce distinct predictive models and features, even if they might achieve similar performance for sex classification.

### Better classification performance is associated with stronger inter-subject synchrony

Having shown the variances in brain states and performance of sex classification across different movie stimuli, we next investigated how the brain state influenced sex classification. We averaged the inter-subject synchrony values over all ROIs for each movie clip to represent the overall brain state and correlated them with sex classification accuracies (Fig. 6A). We found that the overall inter-subject synchrony level was positively correlated with classification performance (r = 0.70, p = 0.007). This result suggests that a movie clip evoking stronger inter-subject synchrony across the whole brain tended to result in a better sex classification performance.

**Fig. 6:**
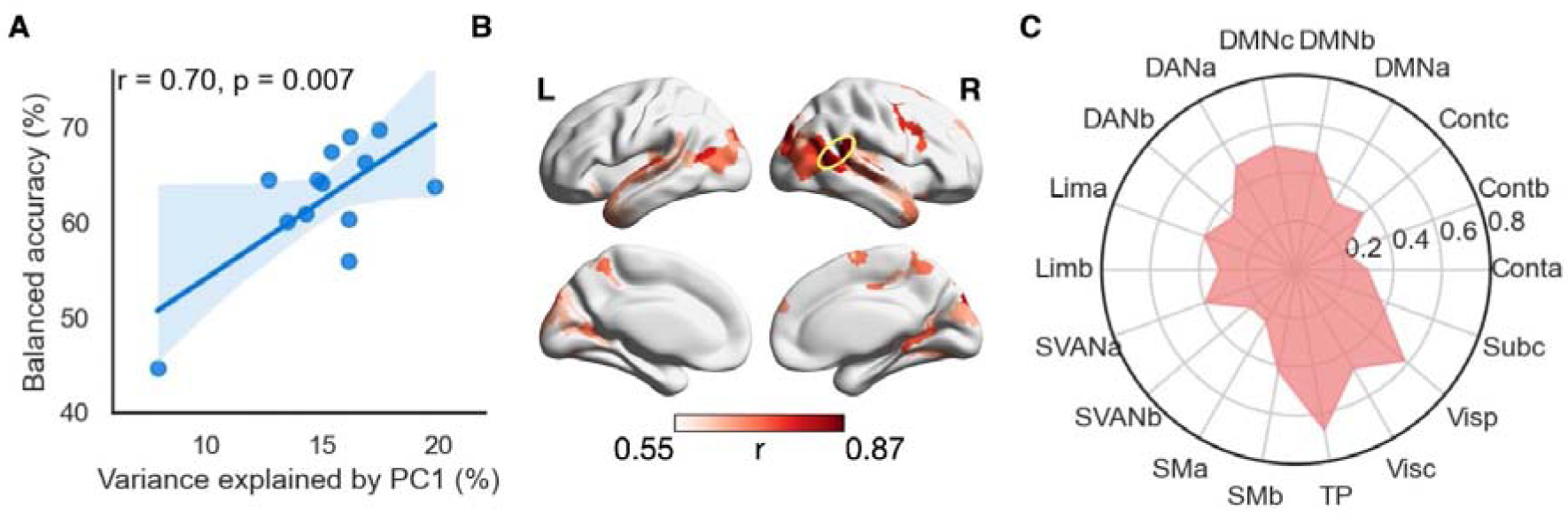
Relationship between classification performance and inter-subject synchrony. **A**) Scatter plot of the relationship (Pearson’s r) between sex classification accuracies and inter-subject synchrony across clips. Each dot represents a movie clip. The line represents the linear regression model fit of the data, with the 95% confidence interval marked by the shade. **B**) ROIs showing a significant (uncorrected p<0.05) correlation between inter-subject synchrony and classification accuracies. Three neighbouring ROIs surviving the correction for multiple comparisons are marked in a yellow circle. Larger correlation values are indicated by darker colours. **C**) Correlations of individual ROIs between inter-subject synchrony and classification accuracies averaged by network.

To examine how inter-subject synchrony influenced feature importance, we correlated the synchrony values with the feature importances across all ROIs for each movie clip (Supplementary Fig. 4A). We did not observe significant correlations between these two measures for most movie clips, except for “hotel” and “dreary” (hotel: r = 0.16, dreary: r = 0.19, both corrected p < 0.05). This result suggests that a brain region showing high inter-subject synchrony is not necessarily an important feature for sex classification. However, we found that the similarity between movie clips in the topography of inter-subject synchrony was significantly correlated with both their similarity in feature importance maps (r = 0.34, p = 0.003; Supplementary Fig. 4B) and cross-movie generalisability (r = 0.43, p <0.001; Supplementary Fig. 4C). These results suggest that movie clips evoking similar spatial patterns of inter-subject synchrony tended to yield similar predictive features and models.

To better understand where in the brain the level of inter-subject synchrony was relevant to sex classification, we computed the correlation between inter-subject synchrony and classification performance for each ROI separately (Fig. 6B). We found that better classification performance was significantly associated with higher inter-subject synchrony levels in three ROIs in the right TPJ (all r = 0.87, FDR corrected p<0.05), a key region for social cognition (Van Overwalle, 2009) and attention maintenance (Langner & Eickhoff, 2013). Fig. 6C shows the correlations grouped by network (Yeo et al., 2011). We found that inter-subject synchrony of the temporal-parietal, visual and default mode networks was more strongly correlated to classification performance than the other networks.

Taken together, these results suggest that better sex classification performance was associated with higher overall inter-subject synchrony across the whole brain. Furthermore, both the magnitude and location of inter-subject synchrony had an impact on sex classification performance.

### Influence of movie features on inter-subject synchrony and classification performance

In the previous sections, we have shown that different movie clips can elicit distinct brain states and result in different utility for sex classification. Then the question arises: what features of these movie stimuli caused these differences? To address this question, we extracted and analysed various features of the movie stimuli that characterise both low- and high-level properties of the movies.

Given that all movie stimuli used in this study were clips either from independent films (CC) or Hollywood films (HO), we first investigated whether sex classification performance differed between the two movie types. Specifically, “inception”, “social net”, “ocean 11”, “home alone”, “brockovich” and “star wars” belong to Hollywood films, and the others belong to independent films (Supplementary Table 1). We compared the classification accuracies averaged over movie clips within each film type between the two film types across all CV folds by using a corrected resampled t-test (Fig. 7A). We found that Hollywood movie clips significantly outperformed independent movie clips in sex classification (CC: 59.1±14.2%; HO: 66.5±13.0%; p = 0.00036). Moreover, we investigated the difference in inter-subject synchrony between the two film types (Fig. 7B). We averaged the inter-subject synchrony values over clips within each film group for all ROIs and compared them between the two groups using a permutation test for paired sample (5000 iterations). Similarly, Hollywood movie clips evoked significantly higher level of inter-subject synchrony than independent movie clips (CC: 13.9±9.6%; HO: 16.4±11.3%; p = 0.0004).

**Fig. 7:**
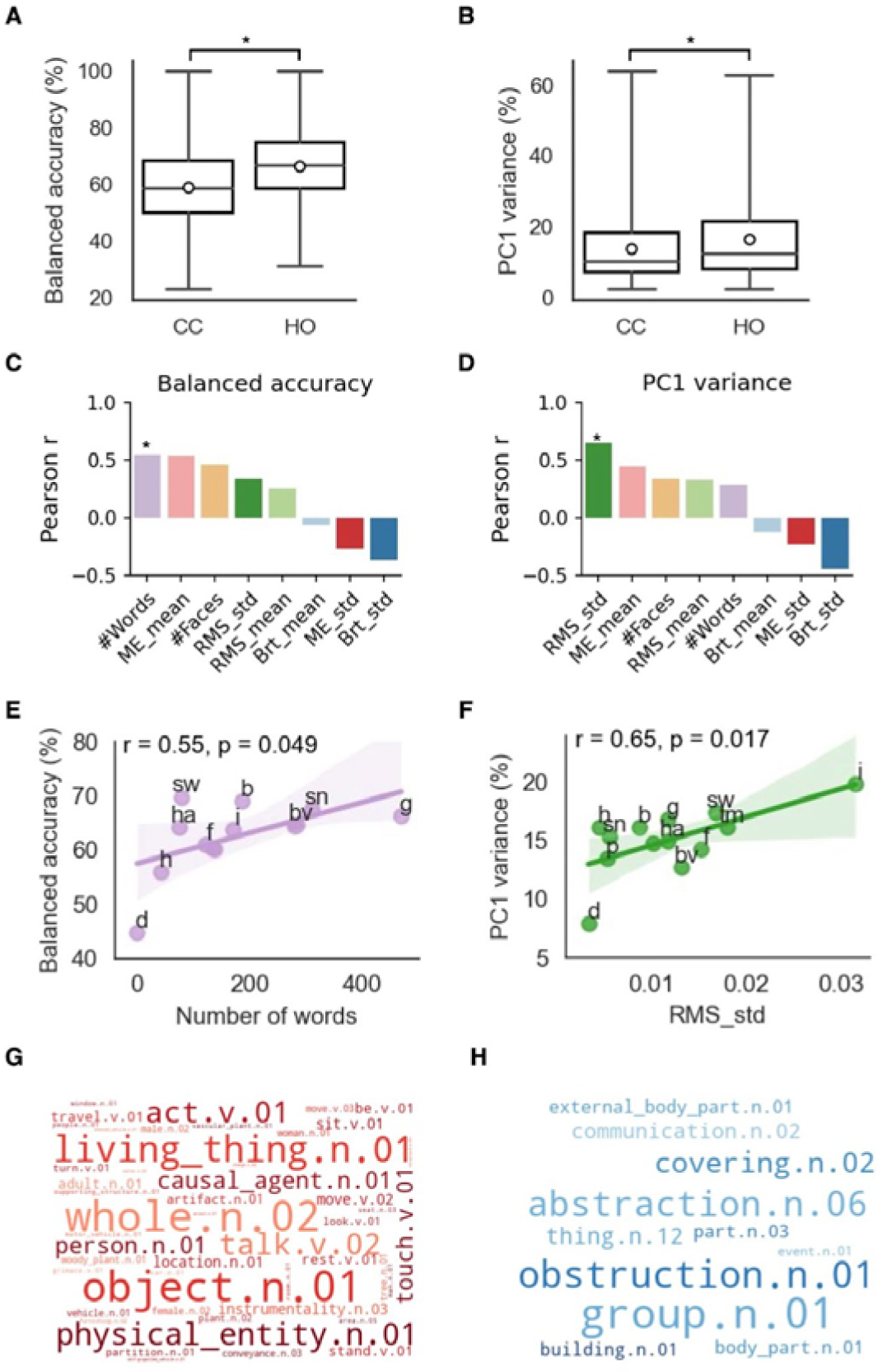
Multi-level features of movie stimuli in relation to sex classification performance and inter-subject synchrony. Comparison between independent (CC) and Hollywood (HO) movie clips in sex classification performance (**A**) and in inter-subject synchrony (**B**). Significance was evaluated by a corrected resampled paired t-test and by a permutation test (5000 iterations) in (**A**) and (**B**), respectively. * denotes p<0.05. Boxplot convention is consistent with that used in Fig. 4A. (**C**) and (**D**) show the correlations (Pearson’s r) between each feature and sex classification performance and the correlations between each feature and inter-subject synchrony, respectively. * denotes p<0.05. **E**) Scatter plot of the relationship (Pearson’s r) between sex classification accuracies and the number of spoken words (#Words) in movie clips. **F**) Scatter plot of the relationship (Pearson’s r) between inter-subject synchrony and the standard deviation across time in RMS of audio (RMS_std). For (**E**) and (**F**), the plot convention is consistent with that used in Fig. 4C. Movie names are annotated in abbreviations (Supplementary Table 1). Word clouds of semantic features showing a positive (**G**) or negative (**H**) correlation coefficient with sex classification performance. The size of each word is proportional to the absolute value of its correlation value.

Next, we investigated several low- and middle-level sensory features of the movie clips. For low-level features, we extracted total motion energy (ME) (Nishimoto et al., 2011), visual brightness (Brt) and auditory loudness (root-mean-square, RMS) at each TR. The mean and standard deviation over all TRs within each movie clip were computed for each feature. For middle-level features, we computed the total number of TRs showing human faces (#TR_faces) and the total number of spoken words (#words) for each movie clip by using automatic feature extraction toolboxes (McNamara et al., 2017) with deep learning based-models for speech recognition (Radford et al., 2022). This resulted in 8 different features altogether covering both visual and auditory aspects of movies. For each feature, we examined its relationships with classification performance and inter-subject synchrony separately across movie clips by using Pearson’s correlation (Fig. 7C and D). We found that classification performance was positively related to the number of spoken words (r = 0.55, p = 0.049; Fig. 7E), suggesting that better sex classification performance was associated with richer speech content in the stimuli. Inter-subject synchrony was positively correlated with the standard deviation of RMS (RMS_std) across TRs (r = 0.65, p = 0.017; Fig. 7F). A high value of RMS_std indicates large variations in loudness across time of a movie clip. Similar patterns were obtained when using partial least squares regression to relate these features to classification accuracy (Supplementary Fig. 5A) and inter-subject synchrony (Supplementary Fig. 5B).

Finally, we explored the influence of high-level semantic features on classification performance and inter-subject synchrony. A total of 859 semantic categories (Huth et al., 2012) were labelled by the HCP for the movie scene at each TR separately. For each movie clip, we computed the appearing frequency of each semantic feature as the number of TRs showing the respective feature divided by the total length of the clip (i.e., 132 TRs). We disregarded features with zero values in over half of the clips and those with the same values across all clips, yielding 70 features. Their relationships with classification performance across all movie clips were examined by using Pearson’s correlation for each feature separately (Supplementary Fig. 5C). We illustrated semantic features that were positively (max r = 0.81) or negatively (min r = -0.42) correlated with classification performance by word clouds separately (Fig. 7G and H). We found that semantic features related to humans, lives and concrete objects (e.g., “living_thing”, “person”, “physical_entity”), actions and social interactions (e.g., ”talk”, ”touch”, ”act”), and story structure (e.g., “causal_agent”) achieved high positive correlations with classification performance (marked in red with large word size). By contrast, words related to abstract concepts (e.g., “group”, “obstruction” and “abstraction”) achieved negative correlations (marked in blue). We repeated the same analysis for the overall inter-subject synchrony and observed very similar patterns of the semantic features (min r = - 0.26; max r = 0.77; Supplementary Fig. 5D-F).

### Control analyses

We conducted several control analyses to ensure the robustness of our results to different choices of machine learning settings, length of movie clips and physiological factors. First, we used three alternative classifiers, i.e., a SVM classifier with a linear kernel (svm_linear), a ridge classifier (ridge) and a random forest classifier (rf), for sex classification instead of the SVM classifier with a RBF kernel (svm_rbf). All classifiers achieved very similar patterns of performance across movie clips to the svm_rbf classifier except the random forest classifier (svm_linear: r = 0.98, ridge: r = 0.94, rf: r = 0.49; Supplementary Fig. 6).

Second, we tested different choices of feature spaces for sex classification. In addition to PC1 loadings, we used PC2 loadings of all ROIs and the concatenation of PC1 and PC2 loadings of all ROIs as features separately, yielding 436 and 872 features respectively. We found that, on average, classification performance was higher when using PC1 loadings (62.5±6.9%) than using PC2 loadings (61.8±6.3%; Supplementary Fig. 7). Moreover, although classification performance was better when using loadings of both PCs together (64.9±6.7%) than using PC1 loadings alone, the overall pattern across clips remained highly similar between the two conditions (r = 0.95). Furthermore, we tested three different values of threshold (i.e., 3%, 5%, 8%) to preserve only the features for which the variance explained by the PC exceeded the thresholds, a procedure that may improve phenotype prediction performance (Li et al., 2023). Again, classification performance across movie clips remained highly similar to that without feature selection (all r ≥ 0.94; Supplementary Fig. 8). These results suggest that our results in the main analysis are largely robust to various machine learning settings.

Third, we explored the influence of the length of movie clips on classification performance. Instead of truncating all clips to 132 TRs, we conducted sex classification by incrementally increasing data length with a short time step (30 TRs, i.e., 30 s; Supplementary Fig. 9A) and a long time step (120 TRs, i.e., 2 mins; Supplementary Fig. 9B) separately. For both analyses, we observed a consistent pattern that classification performance in general increased as data length increased, with the highest accuracy achieving 72.2% (by “social net” with a length of 240 TRs). More importantly, consistent with our results in the main analysis, we observed large variations in classification performance across movie stimuli regardless of whether we used short (<2 mins) or long (up to 10 min) movie clips. These results further confirm that the specific content, apart from the length, of movie stimuli is an important factor to consider for studies of phenotype prediction on naturalistic data.

Finally, we examined whether our results were influenced by physiological signals, specifically head motion and eye blink rate. They were computed as mean framewise displacement (FD) across TRs and frequency of blinks, respectively, for each subject and then averaged over all subjects within each clip. For head motion, we observed that movie clips presented later within each run incited stronger head motion relative to those presented earlier (Supplementary Fig. 10A). Such increase of head motion over time may reflect subject discomfort or fatigue due to long scan duration. Notably, the strength of head motion showed no significant correlation with classification performance (r = 0.18, p = 0.56; Supplementary Fig. 10B) or inter-subject synchrony of brain activity (r = -0.29, p = 0.34; Supplementary Fig. 10C), suggesting the robustness of our results to head motion. Blink rate was not correlated with inter-subject synchrony (r = -0.24, p = 0.42; Supplementary Fig. 11A) but negatively correlated with classification performance (r = -0.55, p = 0.0496; Supplementary Fig. 11B), with no significant differences in blink rate found between the two sexes for all movie clips (all p > 0.05 after FDR correction; Supplementary Fig. 11C). Blink rate has been thought to be associated with vigilance and attention allocation (Demiral et al., 2023; Maffei & Angrilli, 2019), thus potentially influencing the utility of brain signals for phenotype prediction and providing complementary information for stimulus selection.

## Discussion

By comparing 13 different movie clips, we showed that different stimuli evoked distinct levels of inter-subject synchrony of brain activity. Individual brain activity was generally sensitive to the modulation of the stimulus, but the degree of sensitivity varied across the whole brain. By conducting sex classification based on brain activity evoked during watching the movie clips, we further established differences in the utility for phenotype prediction across the movie stimuli. They not only varied in classification performance but also revealed distinct representations of the same underlying brain-phenotype relationship. Critically, we found that better classification performance was associated with higher inter-subject synchrony. Finally, by examining a variety of features of the stimuli at different levels, we found that movie clips with strong social content and cohesive narratives, often from Hollywood films, enhanced both inter-subject synchrony and sex classification performance.

### Different movie stimuli uniquely modulate brain states

Does the brain state change across different movie stimuli? At the group level, we found that most movie clips exhibited a consistent topography of inter-subject synchrony, with visual, auditory and their association cortices showing higher inter-subject synchrony than the other brain regions (Hasson et al., 2004). Low-level sensory brain regions are believed to closely track the rapidly changing sensory input of the movie stimulus and respond more consistently across subjects, compared to those higher-level brain regions (Hasson, Yang, et al., 2008). However, the movie clips evoked remarkably different levels of inter-subject synchrony, which cannot be explained by their differences in the elicited physiological signals (head motion and eye blink rate). In particular, the movie clip “dreary”, which exclusively features natural scenery without any human images or speech, achieved a remarkably lower inter-subject synchrony level than the other clips. Inter-subject synchrony is often used to indicate how much a stimulus can entrain subjects’ brain activity (Abrams et al., 2013; Hasson, Landesman, et al., 2008; Nummenmaa et al., 2014; Ohad & Yeshurun, 2023). Movie clips evoking higher inter-subject synchrony may be more engaging to the subjects than those evoking lower inter-subject synchrony, for example, by better captivating subjects’ attention and evoking stronger emotional immersion (Jääskeläinen et al., 2021; Ki et al., 2016). Another possibility is that movie clips evoking higher inter-subject synchrony may leave smaller space for individualised interpretations by reducing ambiguity in movie content (Finn et al., 2018; Nguyen et al., 2019).

At an individual subject level, we found that the sensitivity of individual brain responses to stimuli varied largely across the whole brain. Consistent with previous studies (Gao et al., 2020; Li et al., 2023), the representation of inter-individual differences in brain responses to the same stimulus (operationalised as individual expression levels of shared responses) was overall stable across different clips in the sensory association regions, such as STG and superior temporal sulcus (STS). These brain regions play important roles in sensory integration and social perception during naturalistic stimulus processing (Hasson, Landesman, et al., 2008; Lahnakoski et al., 2012; Nummenmaa et al., 2012). These findings suggest that these brain regions may mainly reflect stable personal traits in general processing of movie stimuli. By contrast, in the higher-level and primary sensory regions, the way individuals process stimuli is highly sensitive to the specific content of movie stimuli (Chen et al., 2020; Kröll et al., 2023). Therefore, our results show that brain activity during naturalistic conditions reflect both stable, trait-like characteristics of individuals and state-driven aspects that are influenced by the specific movie stimulus (Geerligs et al., 2015).

On the other hand, we found that the representation of individual differences in these “trait-like” brain regions can also vary largely across certain movie clips. For example, “inception” and “dreary” exhibited distinct representations of individual differences from the other movie clips in a brain region in the dorsal attentional network. The fact that these two movie clips evoked the highest and lowest inter-subject synchrony, respectively, suggests that these changes in representations of individual differences may be due to differences in subject engagement and attentiveness (Ohad & Yeshurun, 2023; Young et al., 2010). In addition, we found that the representation of individual differences in a language-related brain region, the left STG, was highly different for movie clips containing little or no speech content and those containing intense speech content (Mesgarani et al., 2014; Shain et al., 2020). Overall, our results confirm differences in brain states across different stimuli and offer initial insights into how changes in movie stimuli can alter representations in individual differences. These results also suggest that complex movie stimuli hold great potential to tune and shape brain activity in a delicate and flexible manner.

### Selection of naturalistic stimuli is important for studies on phenotype prediction

Does the choice of stimulus matter for studies of brain-behaviour relationships with phenotype prediction? We performed three separate analyses to clarify this question. First, we compared the performance for sex classification between the movie clips and observed substantial differences. We note that our results here may provide very conservative lower bounds due to the relatively short length of the movie clips. We demonstrated that such differences across movie clips were not driven by their differences in subjects’ head motion and were robust to different choices of machine learning settings and length of the movie stimulus (ranging from 30 s to 10 min). These findings together confirm that variations in the utility for phenotype prediction across movie stimuli were mainly driven by differences in their specific content and inherent features. Certain movie stimuli may evoke the brain into a state which brings out or amplifies phenotype-relevant brain signals and thus boost prediction performance (Finn et al., 2017; Greene et al., 2018). Consistent with our findings, differences in prediction performance across movie stimuli have been reported for other phenotypes, such as cognitive ability and personality (Finn & Bandettini, 2021; Li et al., 2023).

In the second analysis, we compared predictive features across movie clips. We found that, on average, the left prefrontal and parietal cortices and bilateral temporal cortices contributed most to sex classification. Some of these brain regions have previously been found to be important for sex classification on resting state fMRI data (Weis et al., 2020; Zhang et al., 2018). The brain regions identified in our study are associated with attention, language, emotion, social cognition and memory processing (Barbey et al., 2013; Carter & Huettel, 2013; Narumoto et al., 2001), consistent with the cognitive domains where sex differences have often been reported (Miller & Halpern, 2014; Ritchie et al., 2018). Importantly, the spatial distribution of predictive features was highly different across movie clips even for those with similar classification performance, suggesting that brain systems contributing most to sex classification varied largely across movie clips. This is in accordance with our previous results that, although movie stimuli are in general audiovisual stimuli, they can differentially engage various brain systems and produce different representations of phenotype-relevant signals.

Lastly, we asked whether the learned predictive models can generalise across movie clips. We found that the cross-movie generalisability only achieved a moderate level and highly depended on the specific movie clips being examined. Movie clips having similar predictive features tended to generalise well to each other. In line with our previous results, this again demonstrates that different movie clips may produce distinct predictive models that highlight different aspects of the same brain-phenotype relationship. Moreover, we found that movie clips with higher within-movie classification performance tended to generalise better to the other clips. This suggests that brain activity evoked by these movie clips may more effectively and reliably reflect sex differences in cognitive processes that are common to naturalistic conditions, e.g., emotion or memory processing (Hasson et al., 2015). Such enhancement of phenotype-relevant brain signals may result from better suppression of noise signals, e.g., by improving subjects’ engagement and arousal level (Vanderwal et al., 2015).

Altogether, these results confirm the necessity of a careful selection of naturalistic stimuli. Recent studies have noticed differences in the utility of movie stimuli for studying individual differences and brain-behaviour relationship (Finn & Bandettini, 2021; Gruskin et al., 2020; Li et al., 2023). Our study extends these studies by showing that not only prediction performance but also representations of the brain-phenotype relationship can vary across different stimuli. Furthermore, these findings highlight both challenges and potential in applications of naturalistic conditions for phenotype prediction. Care should be taken when interpreting the learned brain-phenotype relationship with regard to its generalisability across different stimuli. On the other hand, a careful selection of stimuli may allow researchers to better harness the potential of naturalistic conditions for studying brain-behaviour relationships (Eickhoff et al., 2020; Finn et al., 2020). This realisation opens up a lot of possibilities for future studies. For example, future work could consider combining multiple brain states during watching of different movie clips for learning a more reliable and comprehensive representation of brain-phenotype relationships (Tetereva et al., 2021).

### Inter-subject synchrony can be an effective indicator for stimulus selection

What kind of brain state benefits phenotype prediction? For naturalistic studies on shared responses across subjects, manipulating the brain into a state that maximises inter-subject synchrony may be a favourable option. However, when the goal is to study individual differences and phenotype prediction, how much synchrony is desirable remains largely unclear (Finn et al., 2020). In this present study, we found that at least for the phenotype, sex, better predictions were associated with higher overall inter-subject synchrony. Specifically, we found that better sex classification performance was associated with higher synchrony in TPJ and visual, somatosensory, and default mode networks. These brain regions and systems have been found to involve in several high-order functions during naturalistic stimuli processing, such as emotion processing (Nummenmaa et al., 2012), subject engagement and attention (Langner & Eickhoff, 2013; Yeshurun et al., 2021) and episodic memory formation (Hasson, Furman, et al., 2008; Simony et al., 2016), and are also in line with our previous findings on important features for sex classification.

A higher inter-subject synchrony could be a result of stronger activation of certain brain functions. For example, previous studies have observed improved inter-subject synchrony in emotion circuits when subjects perceive stronger emotions from the stimuli (Nummenmaa et al., 2014; Schmälzle et al., 2015). A higher inter-subject synchrony could also indicate better subject engagement and higher attention levels (Ki et al., 2016; Ohad & Yeshurun, 2023). From a methodological perspective, enhancing inter-subject synchrony could increase signal-to-noise ratio by suppressing stimulus-irrelevant signals (e.g., caused by tiredness or mind wandering) while boosting phenotype-relevant signals, thus improving prediction. Moreover, higher inter-subject synchrony may allow for a more reliable and meaningful characterisation of individual differences (i.e., expression levels of shared response) in the TOPF approach (Li et al., 2023). Actually, the TOPF approach belongs to the inter-subject correlation (ISC) family of approaches (Hasson et al., 2004; Nastase et al., 2019; Simony et al., 2016), probably the most widely used family of approaches in naturalistic fMRI studies. As characterisation of individual differences in ISC-based approaches typically all rely on inter-subject synchrony, our findings in this study may also apply to the other ISC-based approaches. Further investigations are needed to explore how inter-subject synchrony influences predictions based on other brain features, such as functional connectivity.

Overall, these results are very intriguing, because one would intuitively expect that higher inter-subject synchrony means less individual variations in brain activity, thereby less useful for studying individual differences and phenotype prediction. In fact, previous studies on functional connectivity have shown that task and naturalistic conditions that make brain states more similar across subjects tend to yield more stable and identifiable individual differences (Finn et al., 2017; Vanderwal et al., 2017). Our results are in line with these studies and provide further evidence that phenotype prediction can be boosted in conditions that make brain states more similar across subjects. Notably, all movie clips used in our study still leave a large room for inter-subject variability, with the shared response accounting for only 20% of the total variance in brain activity across subjects at most. In sum, these findings provide crucial insights into how inter-subject synchrony affects phenotype prediction and underscore the potential of inter-subject synchrony as a powerful indicator for stimulus selection.

### Stimulus features can influence inter-subject synchrony and phenotype prediction

What features make a movie stimulus evoke high inter-subject synchrony and useful for phenotype prediction? We gained several empirical insights into this question by analysing a variety of movie features at different levels. Specifically, we observed that Hollywood movie clips achieved significantly higher inter-subject synchrony compared to independent movie clips. From a cinematic point of view, this result may come from a distinctive feature of Hollywood movies: they often have a high degree of focus on storytelling by using advanced cinematography techniques, professional sound and music designs, and conventional storytelling techniques, such as clear protagonists and antagonists and linear plotlines (Bordwell, 2006; Hasson, et al., 2008; Smith et al., 2012). These techniques may make Hollywood movies easier to understand and appealing to broader audiences than independent movies, resulting in greater inter-subject synchrony. In fact, previous studies have suggested that inter-subject synchrony of brain activity may provide a quantitative, neuroscientific measurement of the effect of films on viewers for film studies (Hasson et al., 2008). From a neuroscience perspective, Hollywood movie clips are more likely to have been seen by subjects before scanning due to their greater popularity than independent movies. A shared background knowledge or memory about a movie may allow the movie to better captivate subjects during scanning (Chen et al., 2017; Jääskeläinen et al., 2008).

Apart from film types, we found that higher inter-subject synchrony was associated with different features of movie stimuli, including larger variations in loudness across time, increased social content and cohesive narratives. Variation in loudness can be caused by many factors of the audio, such as changes in background music, sound effects and emotions of speakers. Consistent with our result, a previous study has shown that energy-related features and acoustic event density of musical stimuli are associated with enhanced inter-subject synchrony (Trost et al., 2015). Stimuli with larger variations in loudness may better captivate subjects’ attention in a bottom-up manner and produce more reliable responses than the stimuli with less dynamic audio (Ki et al., 2016). Social interaction, such as understanding intentions and emotions of other individuals, is a fundamental ability of humans (Adolphs, 2009). Social cognitive processing of naturalistic stimuli has been previously found to enhance inter-subject synchrony in brain systems supporting episodic memory formation (Hasson, Furman, et al., 2008) and emotion processing (Nummenmaa et al., 2012). Moreover, previous studies have found that movies with structured story and cohesive narratives elicit reliable responses in brain regions supporting information integration over longer time scales and improve the attentional level of subjects, evoking stronger inter-subject synchrony than their less-meaningful counterparts (Hasson et al., 2010; Ki et al., 2016; Simony et al., 2016). Overall, these findings support our result in this present study that both low-level sensory features and high-level context and narrative features of a stimulus have an impact on inter-subject synchrony.

Similarly, we found that better sex classification performance was associated with Hollywood movie clips, rich social content, cohesive narrative and, in particular, increased speech content. These results align with previous findings that brain systems supporting language and social processing exhibited prominent sex differences and contributed most to sex classification (Proverbio, 2017; Weis et al., 2020; Whittle et al., 2011). Our study also recapitulates findings of previous studies that movies with strong social content yield reliable representations of individual differences and improved predictions of cognitive ability compared to those with less social content (Gao et al., 2020; Finn & Bandettini, 2021). Moreover, our work extends these findings by offering novel insights into the modulation of social content through the lens of inter-subject synchrony. Taken together, these results suggest exciting opportunities for future studies to better employ complex naturalistic stimuli by customising and optimising their features and content.

### Limitations

One major limitation of this present study concerns the dataset we used. We chose this specific dataset because it contains both a relatively large sample size and a variety of movie clips of different types. Therefore, this dataset allows us to not only conduct machine learning-based phenotype prediction but also statistically analyse the relationship between prediction performance and different choices of movie stimuli. While highly valuable, this dataset may not be the perfect choice to answer our research questions. First, the presentation order of the movie clips was not counterbalanced during data collection, which may influence brain activity due to changes in the tiredness or arousal level of subjects (Tran et al., 2020). Despite this, we rigorously considered several potential confounding variables such as head motion and eye blinking and obtained highly robust results. Second, this dataset contains only relatively short movie clips (less than 5 mins) from a limited selection of film genres. It remains to be seen whether our findings in this study extend to longer movie clips from a broader range of film types. Third, past viewing experiences regarding the movie clips used in the experiment of subjects were not provided in this dataset. Collecting such information may help future studies to better understand differences between movie stimuli and between individual subjects. Moreover, to emphasise the influence of stimulus selection and obtain robust results, we limited phenotype prediction in this study to a pragmatic case, i.e., sex classification. Future work is needed to test whether our findings in this study are specific to sex classification or generalise to other phenotypes (or continuous prediction as opposed to binary classification).

### Conclusions

In summary, we demonstrate that brain activity can be uniquely modulated by different movie stimuli, resulting in distinct brain states, performance for phenotype prediction and predictive models characterising the brain-phenotype relationship. Movie stimuli that elicit stronger inter-subject synchrony of brain activity tend to obtain better phenotype prediction performance. Such stimuli are mostly derived from Hollywood films and featured by rich social content, cohesive narratives and engaging, dynamic audio. Our findings underscore the importance of selecting an appropriate stimulus for studies on individual differences and brain-behaviour relationships using naturalistic conditions. More importantly, they offer future research practical guidance on stimulus selection from both the brain activity and stimulus features perspectives. Our study also holds particular relevance for naturalistic studies on clinical populations, as these populations may be particularly sensitive to stimulus selection (Bolton et al., 2020; Eickhoff et al., 2020; Gruskin et al., 2020). A suitable selection of stimulus may enhance reliability of measurements of their brain activity, thus promoting clinical diagnosis, prediction, and our understanding of the mechanisms underlying neuropsychiatric disorders.

## Methods

### Participants

We considered all available subjects in the HCP study (S1200 release) who participated in movie watching scanning sessions (Van Essen et al., 2013). Six subjects were excluded from further analyses due to not completing all movie watching tasks, yielding 178 subjects (108 females, age = 29.40±3.31) from 90 unique families. The HCP study was approved by the Washington University institutional review board. Informed consent was obtained from all participants.

### Naturalistic stimuli

The HCP study comprised two movie watching scanning sessions, each session containing two separate runs of a length of roughly 15 mins. During each run, three to four short different movie clips (1 - 4 mins in length) were presented with a rest block of 20s as an interval between movie clips. At the end of each run, a repeat validation clip was presented. The validation clip as well as the shortest clip “overcome” (1’03’’) were excluded, resulting in 13 movie clips in our analyses. Each of the 13 movie clips was presented only once during scanning, with the length varying from 2’22’’ to 4’19’’. Movie clips in runs 1 and 3 were from independent (CC) films, whereas those in runs 2 and 4 were from Hollywood (HO) movies. A brief description of the content and length of these movie clips are provided in Supplementary Table 1. All the movie stimuli can be downloaded from the HCP website (https://db.humanconnectome.org/).

### Imaging data acquisition and processing

All fMRI images during movie watching were acquired at a 7T Siemens scanner (TR = 1000 ms, TE = 22.2 ms, resolution = 1.6 mm^3^). We used the fMRI data preprocessed by the standard HCP pipeline, including steps of motion correction, registration to the standard MNI space, high-pass temporal filtering, 24 motion parameters removal and FIX-denoising (Glasser et al., 2013). The preprocessed data were available from the HCP website. No subjects were further excluded based on the exclusion criterion of having a mean framewise displacement (FD) larger than 0.5 mm (Power et al., 2014). Data during rest blocks were discarded, so that only data during watching movie clips were used. We further removed the first 10 volumes of each movie clip to avoid unstable signals. A whole-brain parcellation (Weis et al., 2020), containing 400 cortical (Schaefer et al., 2017) and 36 subcortical ROIs (Fan et al., 2016), was applied to the remaining data. A mean time series over voxels was extracted for each ROI, subject, and movie clip, using the data processing and analysis for brain imaging (DPABI) toolbox (Yan et al., 2016) (http://rfmri.org/dpabi). For a fair comparison across movie clips, all the resulting time series were truncated to 132 TRs, i.e., the length of the shortest clip, “dreary”, by removing the extra TRs at the end of each clip. Different choices of the length were evaluated in “Control analyses”.

### Eye tracking data acquisition and processing

Eye tracking data were provided by the HCP and collected using an Eyelink S1000 system with two different sampling rates (1000 Hz and 500 Hz). In this study, we only analysed data from subjects with a sampling rate of 1000 Hz for a fair comparison. Subjects were further excluded if their data were unavailable for each movie run. Such information can be obtained from the eye tracking data as well as their metadata provided by the HCP. This resulted in a different number of subjects for each movie run for our analysis, varying from 131 to 135 subjects.

### Inter-subject synchrony

To examine the influence of stimulus selection on brain states at a group level, we computed the inter-subject synchrony level of brain activity during watching different movie clips. For each movie clip, we applied a PCA to the z-score normalised fMRI time series across all subjects for each ROI separately. The derived PCs represent the shared responses across subjects. The variance explained by PC1 was used to quantify the inter-subject synchrony of each ROI (Li et al., 2023). This measure of inter-subject synchrony has been shown to be very similar to the widely used subject-averaged inter-subject correlations (Di and Biswal, 2022; Nastase et al., 2019). A greater value of the variance explained by PC1 indicates that subjects were more consistent in their brain activity over time. To quantify the similarity between movie clips of the spatial pattern of inter-subject synchrony, we computed the Pearson’s correlation coefficients of the variance values between each pair of movie clips across all ROIs, resulting in a 13 by 13 correlation matrix.

### Representation of individual differences in evoked brain activity

To examine the influence of stimulus selection on brain activity at an individual subject level, we computed individual expression levels of the shared response to reflect individual differences in evoked brain activity (Finn et al., 2020; Di & Biswal, 2022). Specifically, in each ROI we computed subject-wise loadings of the PC1 of the z-score normalised fMRI time series across subjects, where PC1 represented the shared response as described above (Li et al., 2013). Each PC1 loading represents the expression level of a subject, with a high value indicating greater resemblance of a subject’s brain activity to the shared response. To test whether the pattern of individual expression levels across subjects was stable across movie stimuli, we correlated these PC1 loadings across all subjects between each pair of clips within each ROI, resulting in a 13 by 13 correlation matrix for each ROI. The average correlation value across all pairs of movie clips was then computed within each ROI, with higher values indicating a more stable representation of individual differences across different stimuli.

### Topography-based predictive framework (TOPF) for sex classification

We applied our previously proposed method, TOPF (Li et al., 2023), for sex classification based on subjects’ brain activity patterns evoked during watching each movie clip separately. Specifically, we applied a PCA to the z-score normalised fMRI time series across subjects for each ROI, with the PC loadings reflecting individual expression levels of the shared responses (i.e., PCs). In our main analyses, we focused on PC1 loadings because we have previously shown that PC1 generally better captures the evoked brain activity and its loadings lead to better prediction performance than the later PCs (Li et al., 2023). The PC1 loadings of all ROIs were used as features for sex classification, yielding 436 features for each subject. Other choices of feature sets were tested in “Control analyses”.

We used a support vector machine (SVM) classifier with a radial basis function (RBF) kernel for sex classification in our main analyses due to its robust performance in previous fMRI studies (Weis et al., 2020). Classification performance was evaluated in a nested cross validation (CV) setting. We adopted a 10-fold CV in the outer loop, where in each fold a predictive model was fitted on the training sample and then used to predict sex of subjects in the test sample. We note that features of test subjects were computed separately from those of training subjects to prevent data leakage. While for training subjects the PC1 loadings were used as features, for test subjects the features were computed as the Pearson’s correlation coefficients between their respective fMRI time series and the PC1s learned on the training sample (Li et al., 2023). We also accounted for the family structure of the subjects by including subjects from the same family in either the training or test set for each fold. Regularisation parameters (i.e., “gamma” and “C”) were optimised via a grid search with inner 5-fold CVs. Classification performance was measured by balanced accuracy to avoid overestimation caused by class imbalance. The above procedure was repeated 10 times with different splits of the sample and the final performance score was computed as the balanced accuracy averaged across the 10 repetitions. The whole procedure was performed in Python based on the julearn packages (https://juaml.github.io/julearn/main/index.html)(Hamdan et al., 2023).

### Significance of predictions and model comparison

We applied a permutation test to evaluate whether classification performance achieved significantly above the chance level for each movie clip. We randomly shuffled the labels (i.e., sex) across subjects 1000 times and for each permutation we rerun the classification pipeline as was described above and computed the average classification accuracy across all CV folds. A p-value was then calculated based on a null distribution constructed by these accuracy values obtained on permuted data. The p-value was defined as the proportion of permutations where the value was higher or equal to the original accuracy value that was derived on the non-permuted data. To test whether the difference in classification performance between two movie clips was significant, we employed corrected resampled paired t-tests to the accuracies across all 100 CV folds for each pair of movie clips (Nadeau & Bengio, 2003). This approach accounted for the fact that CV folds were not independent. The p-values were then corrected for multiple comparisons using FDR.

### Permutation feature importance

We used permutation feature importance to evaluate the importance of each feature for sex classification (Breiman, 2001). Given a model, we randomly shuffled the values of a feature across subjects in the test data. The feature importance was defined as the decrease in the balanced accuracy score when applying the model on the shuffled data. For each model and each feature, we shuffled the data 1000 times and calculated the mean feature importance over all iterations. We then averaged the permutation feature importance over all models within each movie clip for each feature. Features with a positive average importance were identified as an important feature. Calculation of permutation feature importance was conducted by using the “PermutationImportance” function in the ELI5 python package (https://github.com/eli5-org/eli5). To quantify the similarity in feature importance between movie clips, we correlated the importance values between each pair of movie clips across all ROIs.

### Cross-movie generalisability of predictive models

To investigate the generalisability of predictive models across movie clips, we evaluated cross-movie classification performance for each pair of clips based on the models and features obtained in the previous analysis. Specifically, given a pair of clips A and B, in each fold, we applied the model previously fitted on data of clip A to predict sex of subjects in the corresponding test sample based on data of clip B. To improve the comparability across clips, the sign of each PC, as well as of the features, was flipped if the maximum of the corresponding PC loadings was not equal to the maximum of the absolute values of the loadings. Similarly, the cross-movie classification performance was measured by balanced accuracy, which was computed for each fold and then averaged over all folds of all repetitions. Note that the generalisability from clip A to clip B is normally different from that from clip B to clip A.

### Relationship between inter-subject synchrony and sex classification performance

To examine the influence of inter-subject synchrony on classification performance, we first computed the average inter-subject synchrony value across all ROIs for each clip and correlated them with classification accuracies across movie clips using the Pearson’s correlation. Next, we tested the relationship between inter-subject synchrony and feature importance. For each movie clip, we computed the Pearson’s correlation coefficient between the inter-subject synchrony and the permutation feature importance values across all ROIs. Furthermore, we tested how similarity in the topography of inter-subject synchrony between movie clips was related to their similarity in predictive features and models. We correlated the similarity values between movie clips in inter-subject synchrony maps (Fig. 2C) with their similarity values in feature importance maps (Fig. 5C) and cross-movie generalisability (Fig. 5D), separately. Note that for cross-movie generalisability, we averaged the values obtained from two directions for each pair of movie clips to facilitate comparison. Finally, we examined the correlation between inter-subject synchrony and classification performance for each ROI separately. Multiple comparisons were corrected using FDR.

### Comparison between clips from Hollywood films and independent films

We tested whether classification performance and inter-subject synchrony differed between the two types of movie clips (i.e., 7 Hollywood vs. 6 independent films). Specifically, we averaged the balanced accuracy across all clips within each film group for each CV fold separately, yielding 100 mean accuracy values for each group. A corrected resampled t-test was used to examine the difference between the two groups. For inter-subject synchrony, we averaged the variance explained by PC1 across clips within each film group for each ROI separately, yielding 436 values for each group. A permutation test with 5000 iterations was used to examine the statistical significance of the differences between the two groups in inter-subject synchrony.

### Analyses of features of movie stimuli

A variety of features were extracted for each movie clip. Specifically, for each TR, we computed total motion energy (Huth et al., 2012), as the mean over all 4025 channels (provided by the HCP) (Nishimoto et al., 2011). We also extracted visual brightness and auditory loudness (the root mean square of audio signals, RMS) at each TR. Then for each of the three measures, we computed the mean and standard deviation (std) of the values across TRs within each clip, yielding six (four visual and two auditory) low-level features for each clip. We extracted two middle-level features for each clip, i.e., the total number of TRs when human faces were shown (#TR_faces) and the total number of words presented in the audio (#words). #TR_faces, as well as brightness and loudness, were extracted automatically by using a python-based feature extraction toolbox, Pliers (McNamara et al., 2017). For the computation of #words, we used a deep learning based-model for speech recognition provided by Whisper (Radford et al., 2022) to automatically transcribe speech in the audio of each movie clip.

High-level features were extracted based on the 859 semantic (WordNet) labels for each image (Huth et al., 2012), provided by the HCP. For each clip, we computed the appearing frequency of each semantic label as the number of TRs showing the label divided by the total number of TRs of the clip (i.e., 132). Labels that had a zero value for more than half of all clips or had the same value for all clips were excluded, resulting in 70 features for further analysis.

Each movie feature was then correlated to classification performance and inter-subject synchrony values separately across all clips. We note that we accounted for a haemodynamic delay of 4 s when analysing all these movie features.

### Control analyses on the effect of different choices of machine learning settings

We tested whether our results were robust to different choices of machine learning settings by using additional classification algorithms and feature spaces. Specifically, instead of the SVM classifier with a RBF kernel (svm_rbf), we used three commonly used classifiers, i.e., a SVM classifier with a linear kernel (svm_linear), a ridge classifier (ridge_clf) and a random forest classifier, for sex classification.

The procedure for the evaluation of classification performance remained the same for all classifiers. Next, instead of using PC1 loadings as features, we tested two additional feature spaces, i.e., loadings of PC2 (the PC that explains the second largest variance across subjects) and a concatenation of both PC1 and PC2 loadings of all ROIs. These two settings resulted in 436 and 872 features respectively. In addition, if a PC explains too little variance across subjects, its loadings may not reflect meaningful individual differences and thus can introduce noise to the features. Therefore, we applied a feature selection step by using three different values of threshold (i.e., 3%, 5%, 8%). Only features for which the variance explained by their corresponding PCs exceeded the given threshold was used for sex classification. For each of these additional sets of features, we evaluated classification performance for each movie clip with the same procedure as used in the main analysis.

### Control analyses on the effect of length of movie clips

To investigate how the choice of movie length would influence sex classification performance, we conducted two analyses with a wide range of clip lengths. First, for each movie clip, we gradually increased data length (N_tr) by steps of 30 TRs (i.e., 30 s) starting from N_tr = 30 until N_tr exceeded the total length of the given movie clip, yielding a maximum of 8 different data lengths. We then reapplied the TOPF approach to extract features (i.e., PC1 loadings) and perform sex classification for each N_tr and each movie clip separately using the same procedure as described before. Second, we conducted a similar analysis but used a longer time scale by concatenating data across movie clips within each run. We started from N_tr = 120 and increased data length by steps of 120 TRs (i.e., 2 mins) until N_tr = 600, resulting in 5 different data lengths. fMRI data were concatenated in the order of the movie clips as they were presented within each run. Similarly, for each run with each N_tr, we re-extracted features and re-performed sex classification.

### Control analyses on the effect of head motion and eye blinking

We investigated how head motion and eye blinking of subjects observed during watching different movie clips influenced classification performance and inter-subject synchrony. For head motion, we computed the mean FD over all TRs within each movie clip and then averaged the values over all subjects, resulting in one value for each clip. Blink rate was quantified as the number of blinks occurred divided by the total number of TRs during watching the given movie clip. Similarly, the blink rate was computed for each subject and then averaged over subjects, resulting in one value for each movie clip. For each measure, we correlated their values with classification performance and the overall inter-subject synchrony values separately over all movie clips.

## Acknowledgement

This work was supported by the European Union’s Horizon 2020 Research and Innovation Programme under grant agreement no. 945539 (HBP SGA3), and the Deutsche Forschungsgemeinschaft (491111487). Data were provided in part by the Human Connectome Project, WU-Minn Consortium (Principal Investigators: David Van Essen and Kamil Ugurbil; 1U54MH091657) funded by the 16 NIH Institutes and Centers that support the NIH Blueprint for Neuroscience Research; and by the McDonnell Center for Systems Neuroscience at Washington University.

## Supplementary Materials for

**Supplementary Fig. 1:**
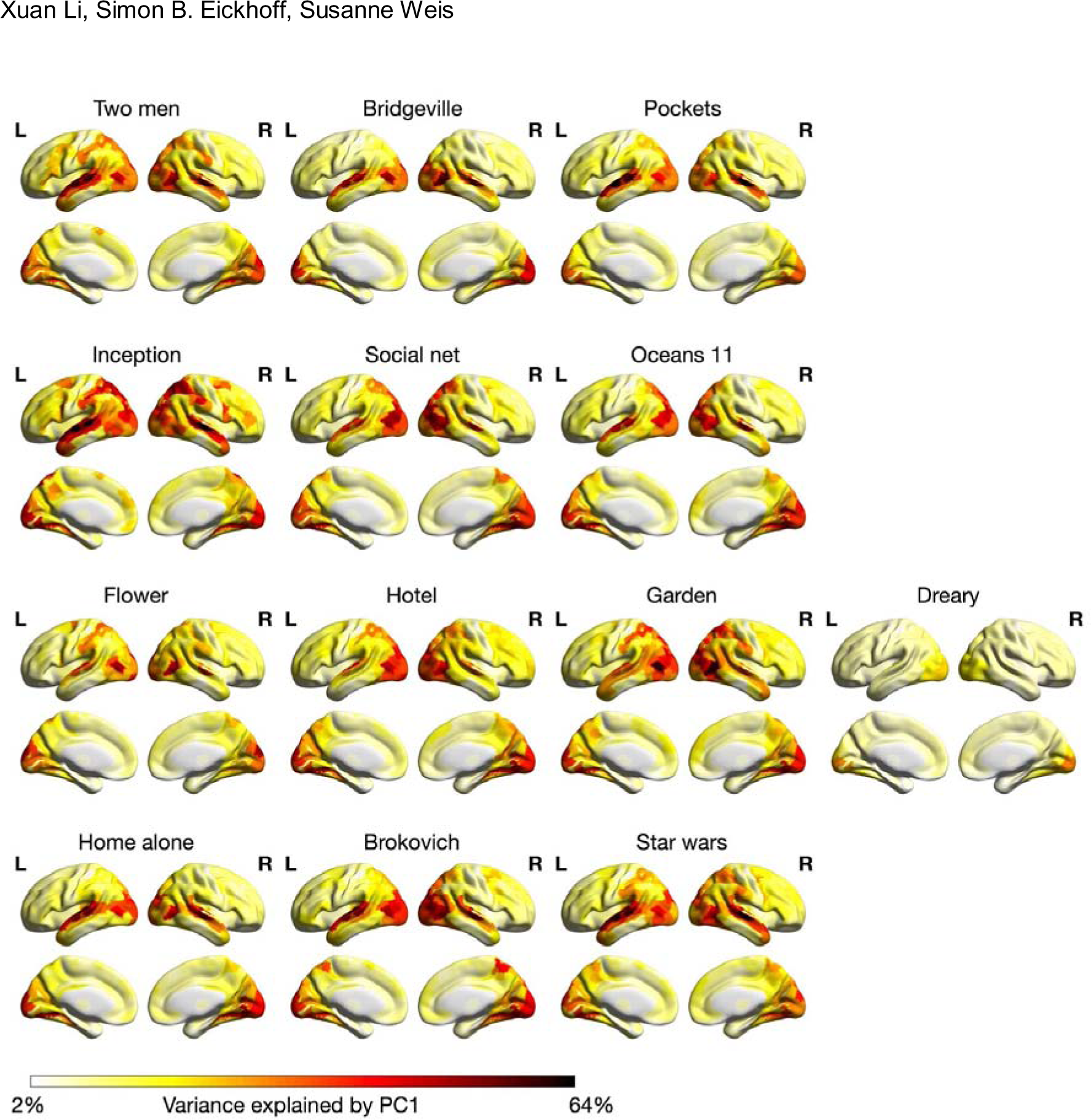
Brain maps of inter-subject synchrony for individual movie clips. Each value represents the inter-subject synchrony of brain activity (quantified as the variance explained by PC1) in a given ROI. High and low inter-subject synchrony values are indicated by dark and light colours, respectively.

**Supplementary Fig. 2:**
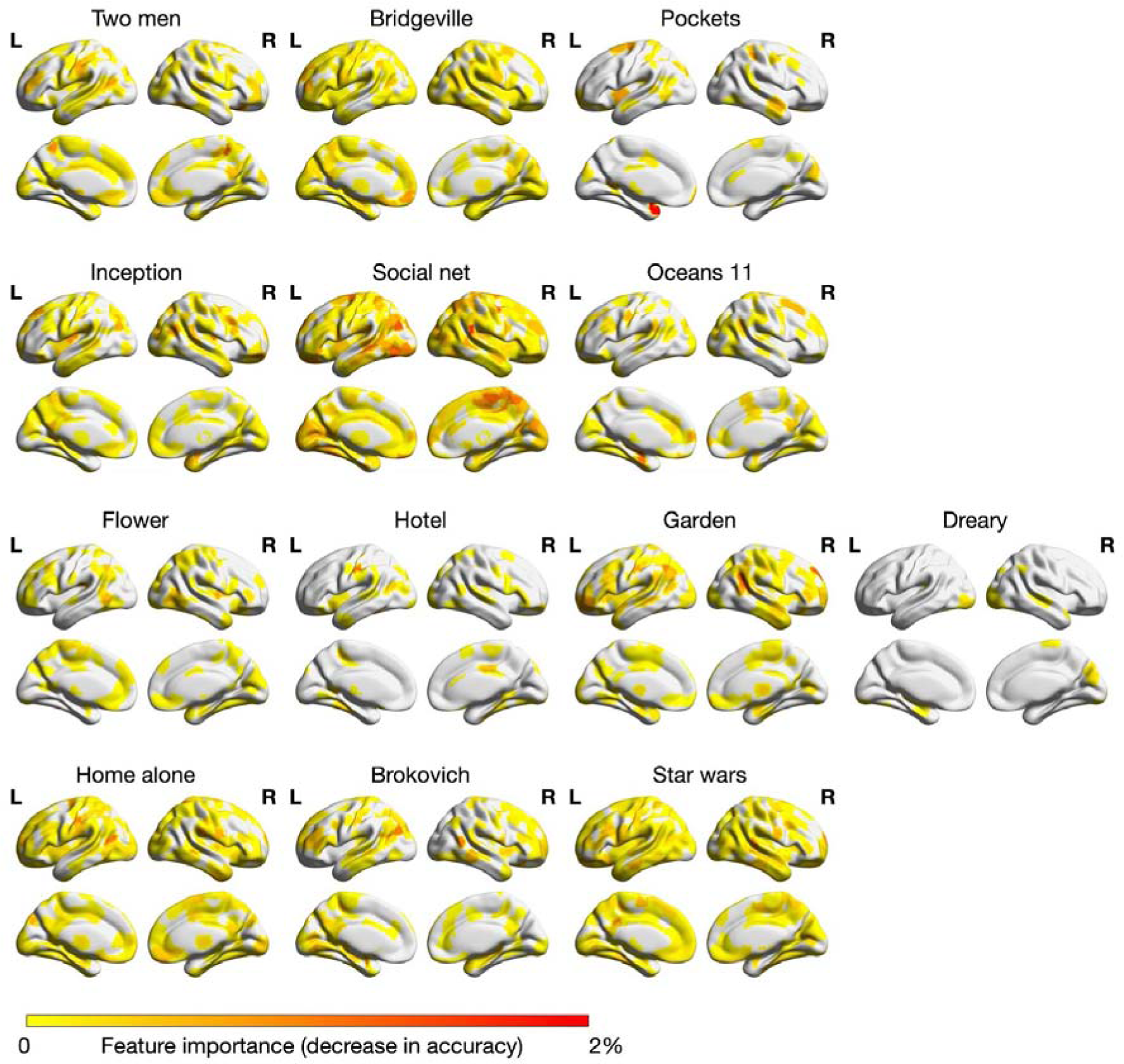
Brain maps of permutation feature importance of all movie clips. Permutation importance of each feature was measured as the decrease in balanced accuracy after shuffling the feature across subjects. Each value in the brain maps represents the average permutation feature importance over 1000 iterations and all predictive models of a given clip. Negative values were set to zeros for illustration. High and low importance values are indicated by the colours red and yellow, respectively.

**Supplementary Fig. 3:**
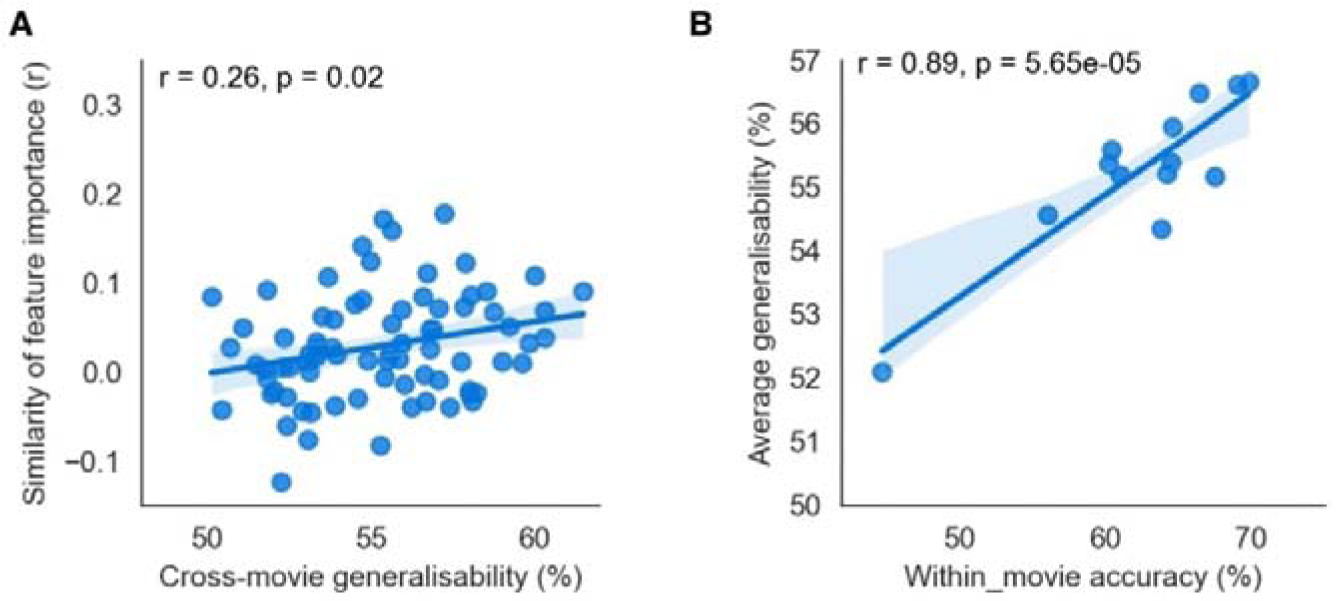
Relationships between feature importance, cross-movie generalisability and within-movie classification performance. **A)**Scatter plot of the relationship (Pearson’s r) between similarity of feature importance (Fig. 5C) and generalisability across movie clips (Fig. 5D). Each dot represents a pair of movie clips. Note that for cross-movie generalisability, we averaged the values obtained from two directions for each pair of movie clips. **B**) Scatter plot of the relationship (Pearson’s r) between average generalisability to the other clips and within-movie classification performance. Each dot represents a movie clip. For both panels, the plot convention is the same as that used in Fig. 6A.

**Supplementary Fig. 4:**
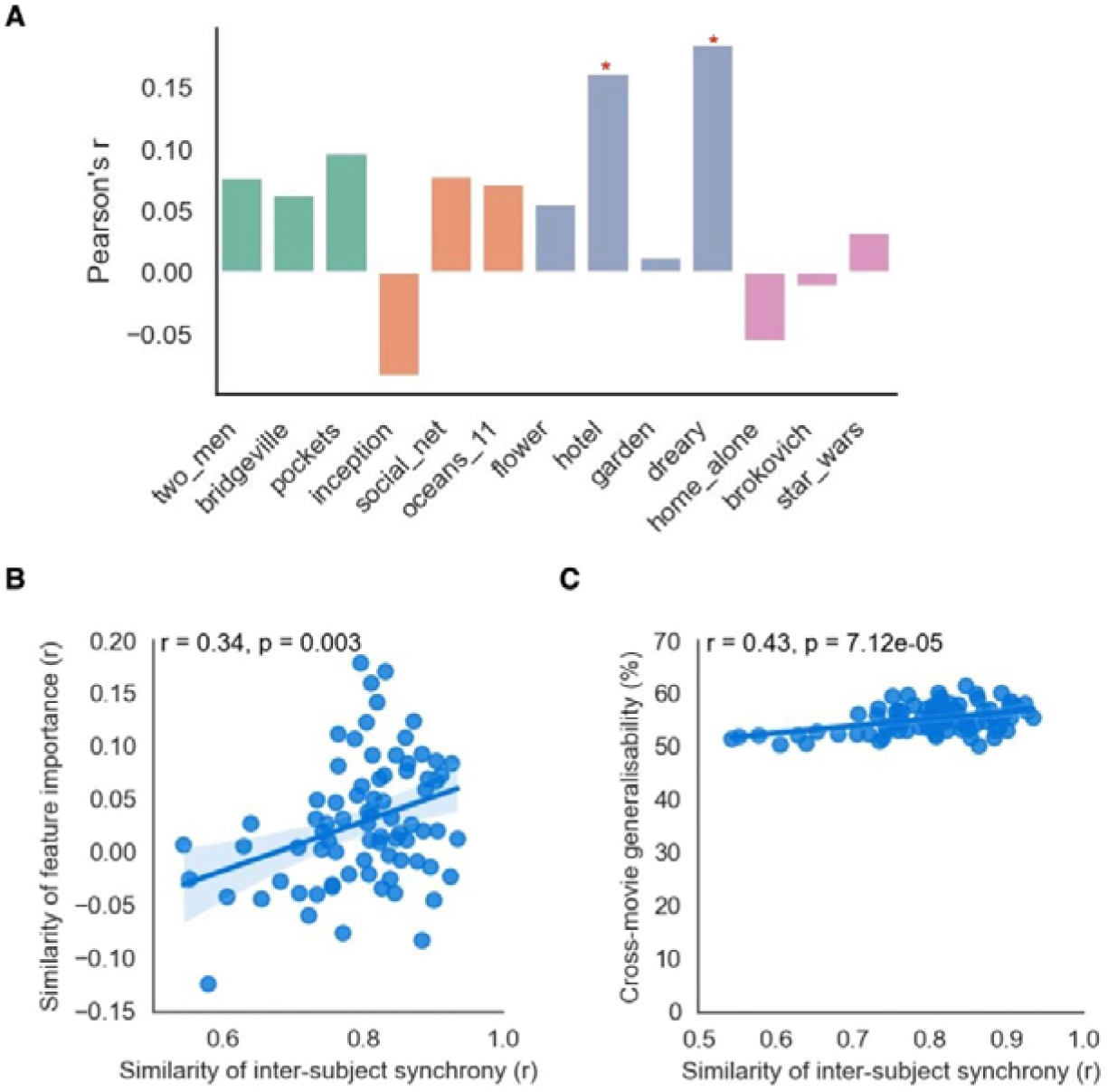
Relationship between inter-subject synchrony and predictive features. **A**) For each movie clip, the correlation (Pearson’s r) between inter-subject synchrony (Supplementary Fig. 1) and feature importance (Supplementary Fig.2) across all ROIs was computed. * denotes p<0.05 (FDR corrected). **B**) Correlation between similarity in inter-subject synchrony (Fig. 2C) and similarity in feature importances across movie clips (Fig. 5C). **C**) Correlation between similarity in inter-subject synchrony (Fig. 2C) and model generalisability across clips (Fig. 5D). For (**B**) and (**C**), each dot represents a pair of movie clips, with the plot convention the same as that used in Fig. 6A. For generalisability, we averaged the values within each clip pair over the two directions.

**Supplementary Fig. 5:**
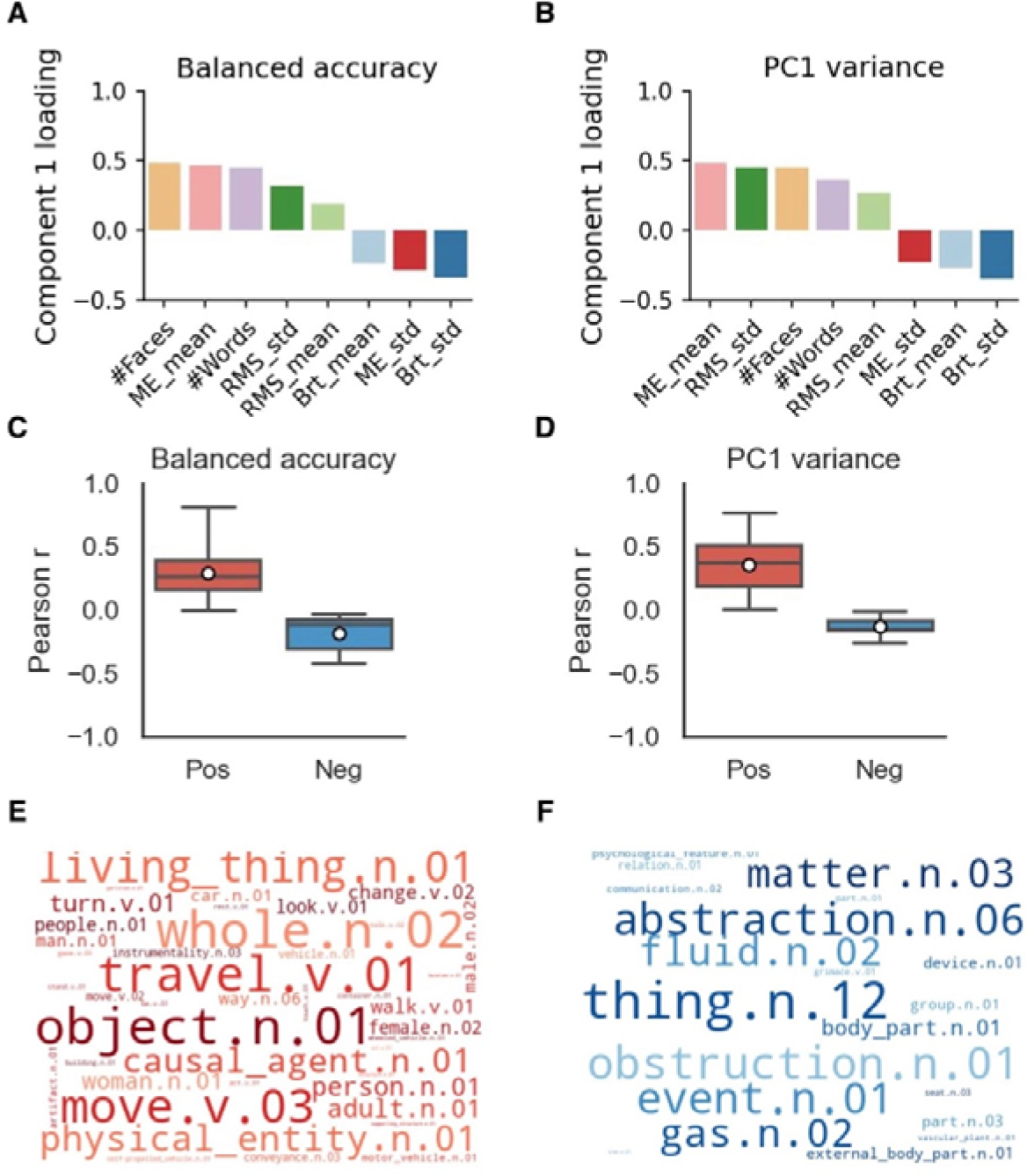
Movie features of movie stimuli in relation to sex classification accuracy and inter-subject synchrony. Relationship between low/middle level movie features and classification accuracy (**A**) as well as inter-subject synchrony (**B**), revealed by partial least squares regression. The y-axis shows the loading of each feature in the first component. The (Pearson’s r) correlation coefficients between the appearing frequency of semantic features and classification accuracy (**C**) and inter-subject synchrony (**D**) across movie clips, grouped by their signs. A positive correlation indicates that better classification accuracy or higher inter-subject synchrony is associated with a higher appearing frequency of a given feature. The boxplot convention is consistent with that used in Fig. 2A. Word clouds of semantic features showing a positive (**E**) or negative (**F**) correlation coefficient with inter-subject synchrony. The size of each word is proportional to the absolute value of its correlation value.

**Supplementary Fig. 6:**
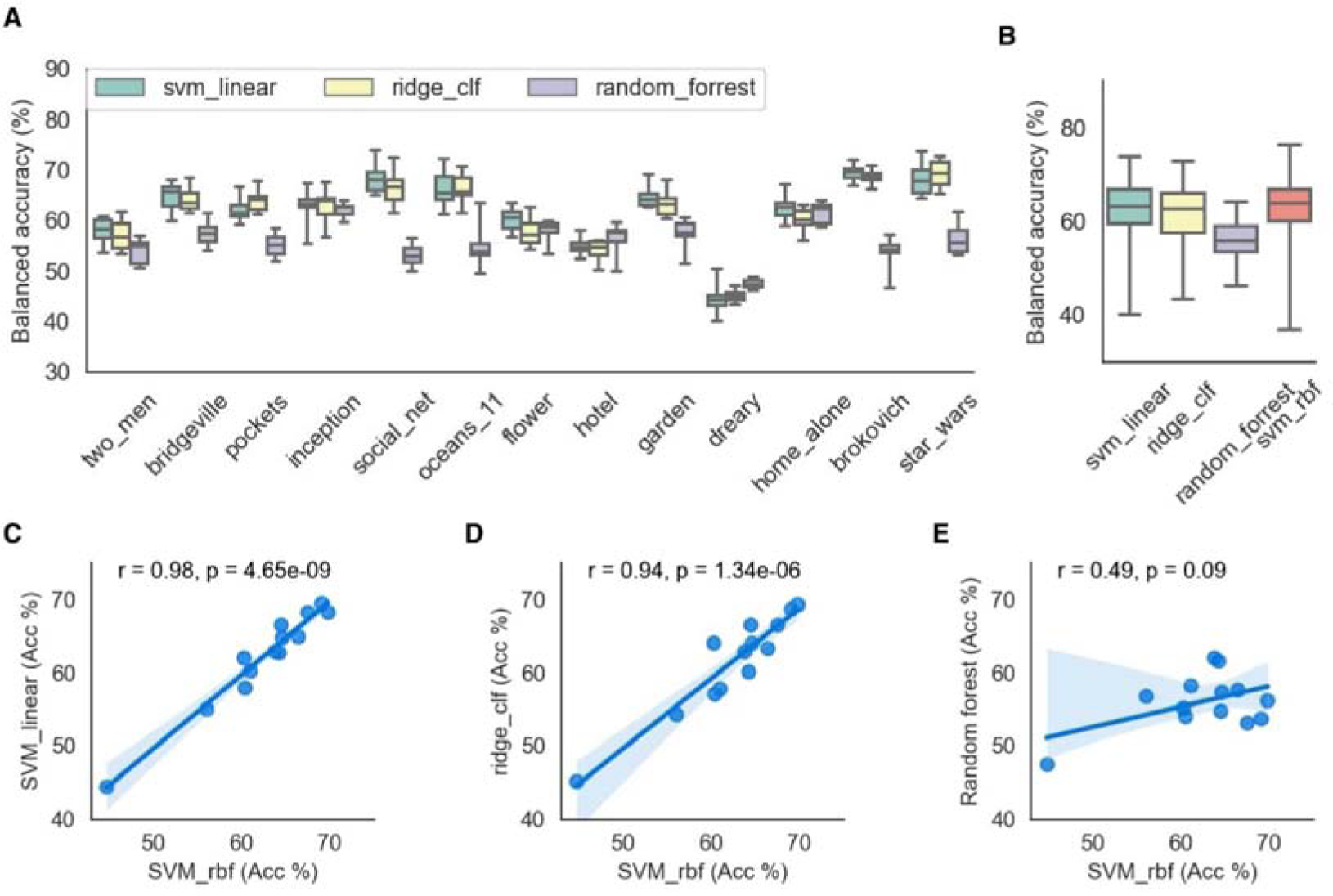
Influence of the choice of algorithms on sex classification performance. **A**) Sex classification performance (balanced accuracy) evaluated in 10 repetitions of 10-fold CV using three different algorithms. The three algorithms are SVM with linear kernel (svm_linear), ridge classifier (ridge_clf) and random forest classifier. **B**) Average accuracy over all movie clips for each algorithm. Similarity (Pearson’s r) of the pattern of balanced accuracy (Acc) over clips between svm_rbf and svm_linear (**C**), ridge_clf (**D**) and random forest (**E**). The plot convention is consistent with those used in Fig. 6A, with each dot representing a movie clip.

**Supplementary Fig. 7:**
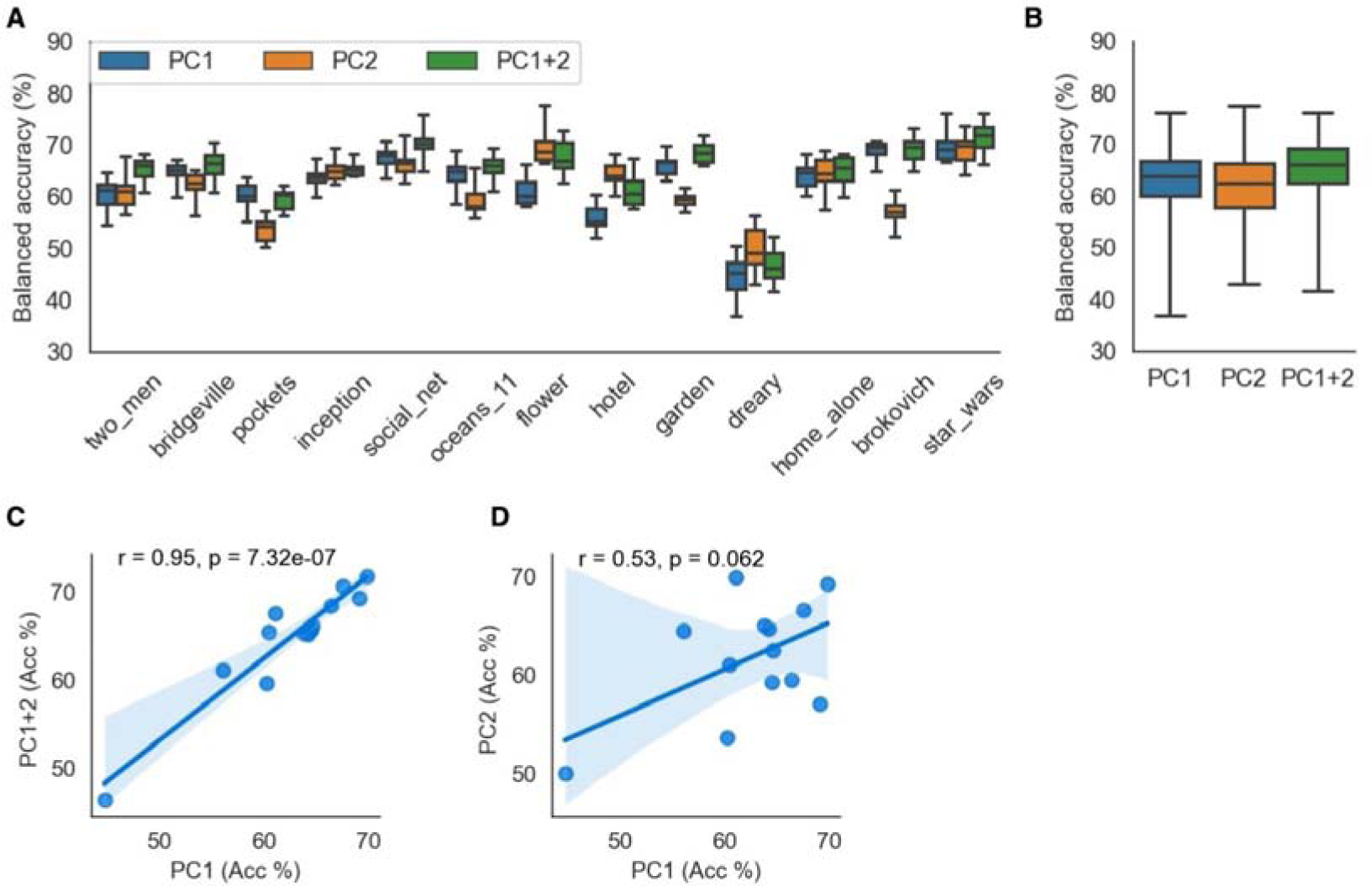
Influence of the choice of TOPF feature sets on sex classification performance. **A**) Sex classification performance (balanced accuracy) evaluated in 10 repetitions of 10-fold CV using three different choices of TOPF features. PC1, PC2 and PC1+2 denote the conditions when using PC1 loadings, PC2 loadings and the concatenation of PC1 and PC2 loadings of all ROIs as features for classification, respectively. **B**) Average accuracy over all movie clips for each condition. Similarity (Pearson’s r) of the pattern of balanced accuracy (Acc) over clips between PC1 and PC1+2 (**C**) and PC2 feature sets (**D**). Each dot represents a movie clip.

**Supplementary Fig. 8:**
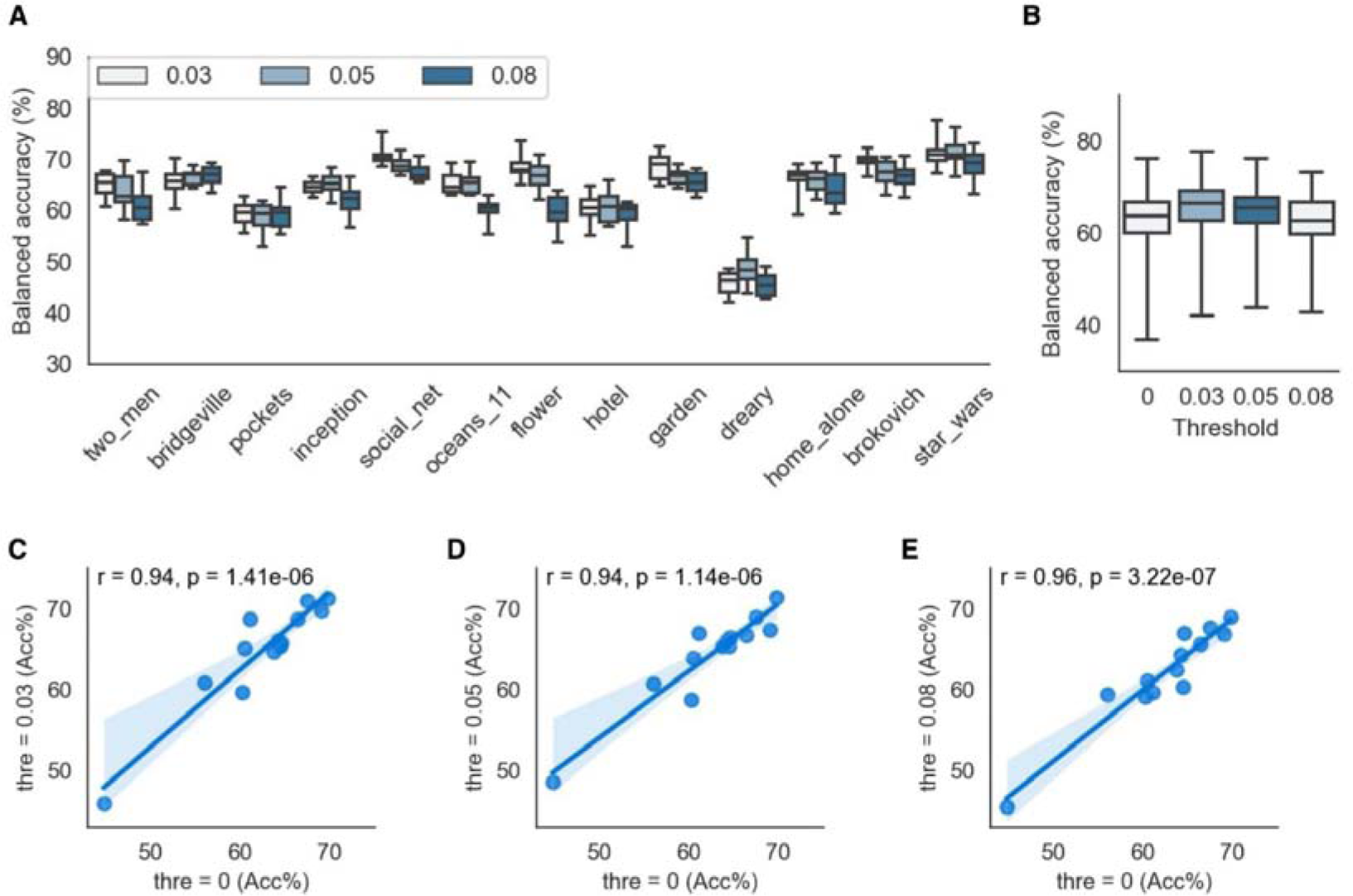
Influence of feature selection on sex classification performance. **A**) Sex classification performance (balanced accuracy) evaluated in 10 repetitions of 10-fold CV using three different values of threshold on the variance explained by PC1 (i.e., 3%, 5% and 8%). **B**) Average accuracy over all movie clips for each condition. The pattern of classification accuracy over clips in our main results without feature selection (threshold = 0%) is highly similar to that using a threshold of 3% (**C**), 5% (**D**) and 8% (**E**).

**Supplementary Fig. 9:**
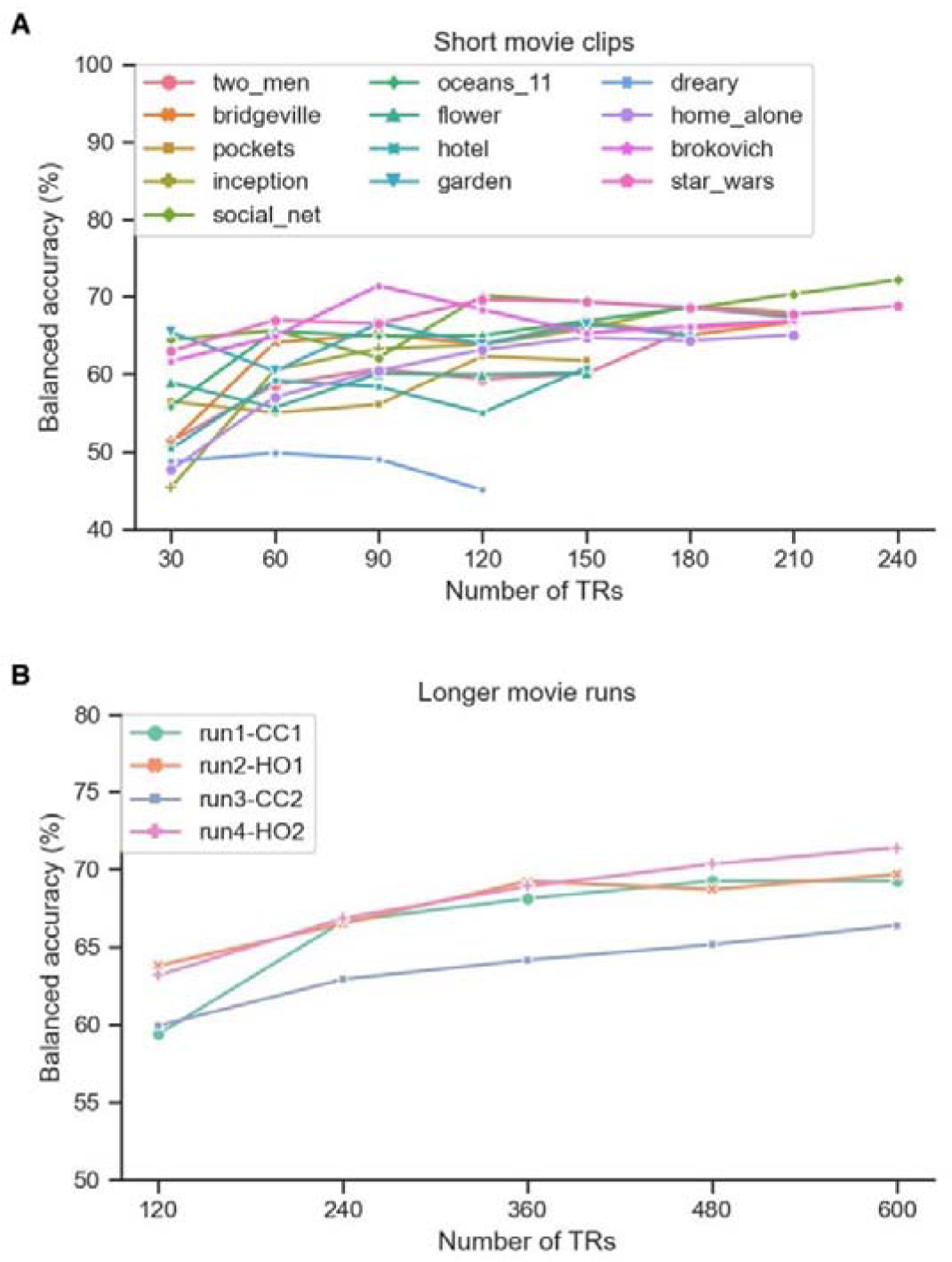
Influence of the length of movie clips on sex classification performance. Sex classification (balanced accuracy) using data with varying short lengths (from 30 s to 240 s with steps of 30 s) (**A**) and with varying longer lengths (from 120 s to 600 s with steps of 120 s) (**B**). Each data point denotes the mean accuracy averaged over all CV folds from all 10 repetitions. TR = 1s.

**Supplementary Fig. 10:**
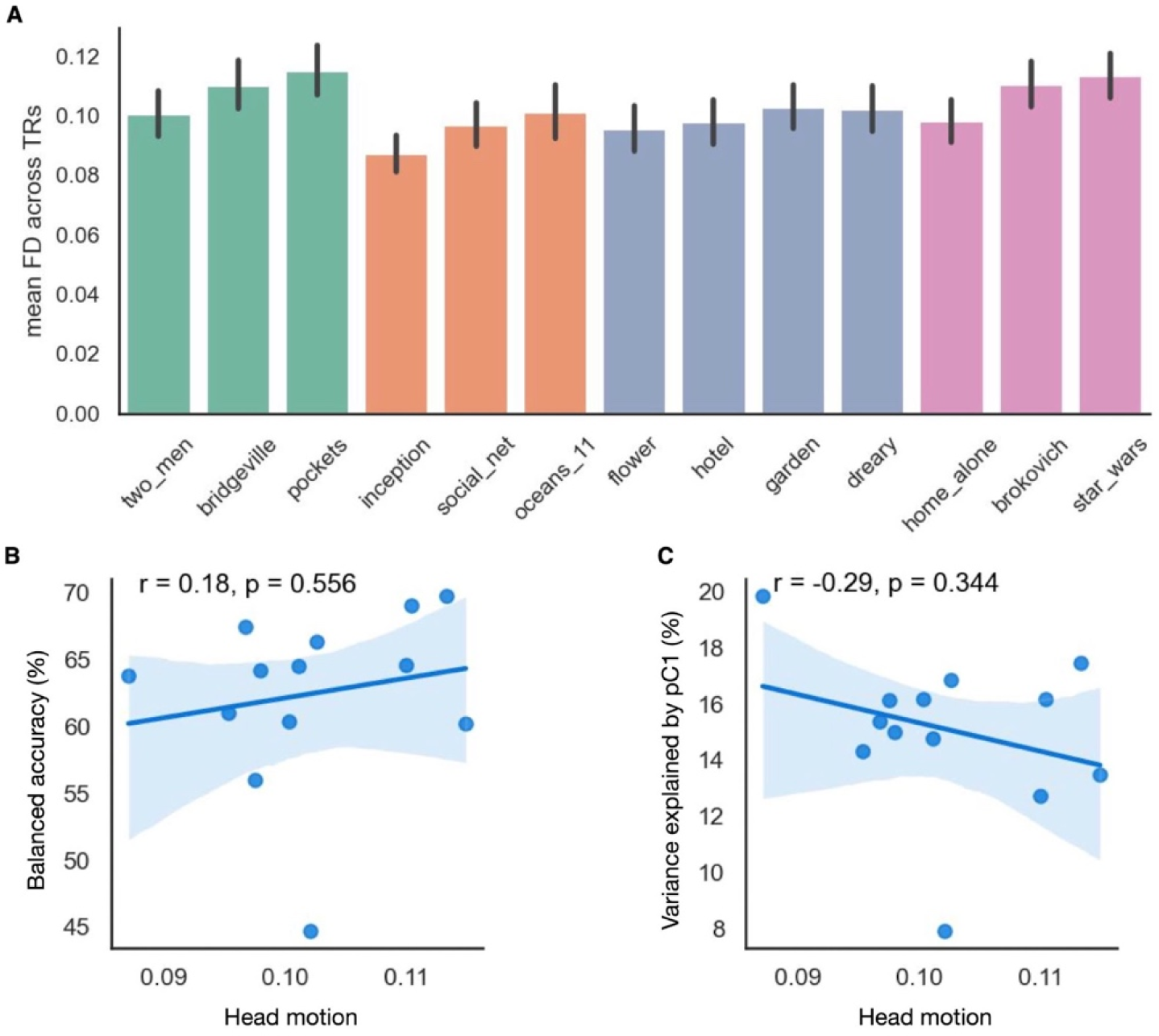
Influence of head motion on sex classification performance and inter-subject synchrony. **A**) Head motion (mean FD across TRs) for each movie clip. The bars indicate the average head motion over all subjects, with the error bar reflecting the standard deviation across subjects. The movie clips are shown in the same order as they were presented within each run from run 1 to run 4. Movie clips belonging to the same run are marked in the same colour. **B**) Scatter plot of the relationship (Pearson’s r) between group-averaged head motion and balanced accuracy. **C**) Scatter plot of the relationship (Pearson’s r) between group-averaged head motion and inter-subject synchrony. Each dot represents a movie clip.

**Supplementary Fig. 11:**
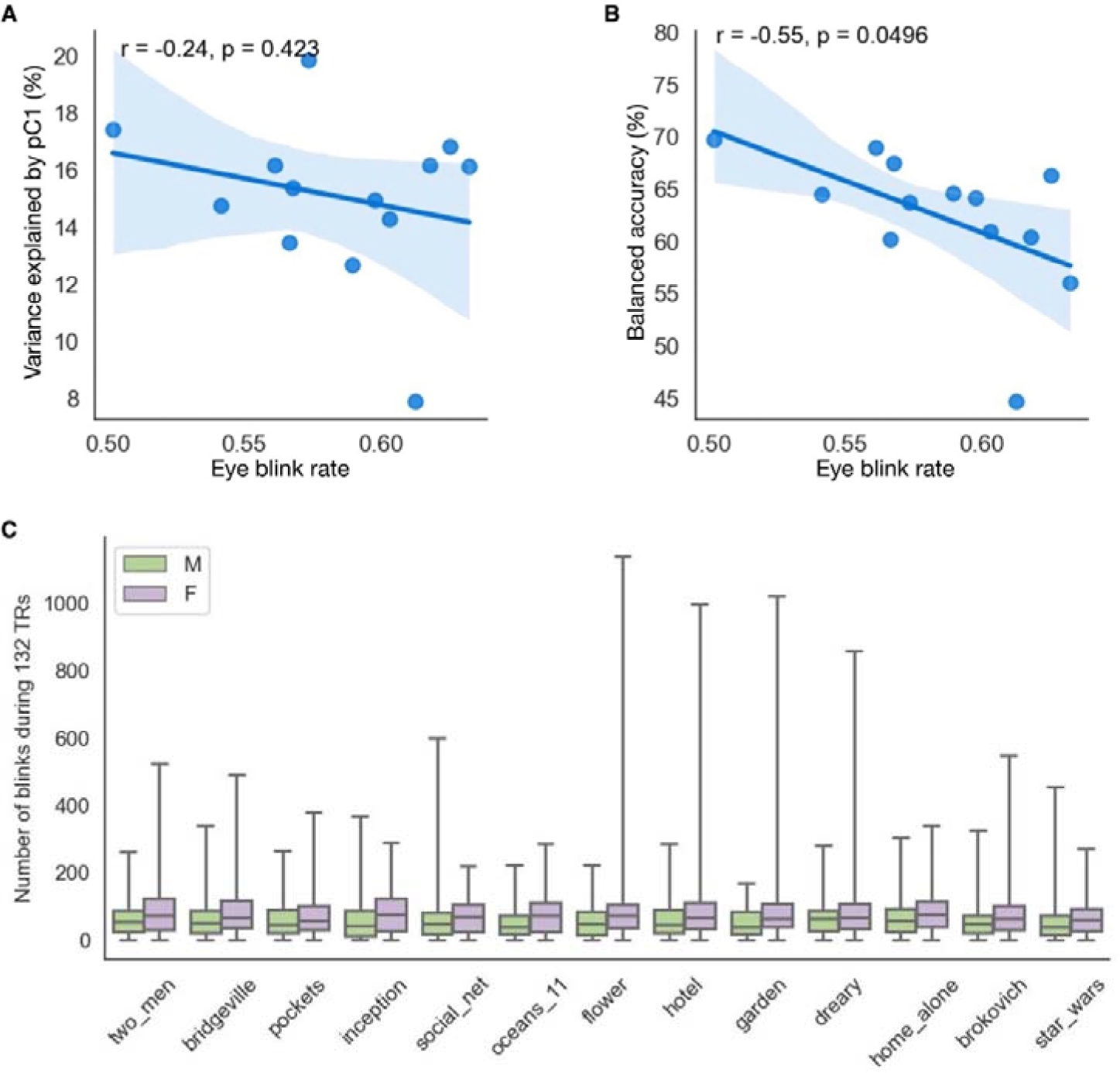
Influence of eye blink rate on sex classification performance and inter-subject synchrony. **A**) Scatter plot of the relationship (Pearson’s r) between group-averaged eye blink rate and inter-subject synchrony. **B**) Scatter plot of the relationship (Pearson’s r) between group-averaged eye blink rate and balanced accuracy. Each dot represents a movie clip. **C**) Comparison of the number of eye blinks between males (M) and females (F) for each movie clip. While on average females consistently showed more eye blinks than males, no movie clips showed a significant difference between the two sexes by t-tests (all p-values > 0.05 after FDR correction).

**Supplementary Table 1:** Summary of the 13 movie clips in HCP used in our analyses. CC and HO denote independent films and Hollywood films, respectively. Abbr. denotes the abbreviation of the name of each clip. Note that in the given durations the first 10 TRs were already excluded for each movie clip.

| Run | type | Short name | Original name | Abbr. | TRs | Duration (min:sec) | Content |
| --- | --- | --- | --- | --- | --- | --- | --- |
| 1 | CC | Two Men | Two Men | tm | 234 | 3:54 | A man is observing another man running in his direction, while narrating his thoughts and thinking whether he should intervene. The narrator speaks accented English and subtitles are given. The last 32 TRs show the credits accompanied by music. |
| 1 | CC | Bridgeville | Welcome to Bridgeville | b | 211 | 3:31 | Inhabitants of a town tell stories about them enjoying living there. The speakers are shown with montages of the town and accompanied by music. |
| 1 | CC | Pockets | Pockets | p | 179 | 2:59 | People show what they keep in their pockets and explain their personal meaning. Clips show close-ups of their items and faces, accompanied by music. The last 9 TRs show credits. |
| 2 | HO | Inception | Inception | i | 217 | 3:37 | A man and a woman walk around at an imaginary location, learning how to interact with their surroundings and its possible dangers. |
| 2 | HO | Social Net | The Social Network | sn | 249 | 4:09 | The clip consists of many short scenes. It shows young Zuckerberg at university at a hearing where he is being accused of breaking rules. The last scene shows Zuckerberg leaving a lecture emotionally agitated. The scenes are re-enacted. |
| 2 | HO | Ocean's 11 | Ocean's Eleven | o | 239 | 3:59 | A man introduces his later accomplices to his plan on how to rob a casino. The men discuss their approach. |
| 3 | CC | Flower | Off the Shelf | f | 170 | 2:50 | A song with montage of a flower and its journey through different places |
| 3 | CC | Hotel | 1212 | h | 174 | 2:54 | A man and a woman are in a hotel room and supernaturally experience the presence of each other but never meet. |
| 3 | CC | Garden | Mrs Meyer's Clean Day | g | 195 | 3:15 | A documentary of a woman presenting her community program teaching people how to grow food. |
| 3 | CC | Dreary | Northwest Passage | d | 132 | 2:12 | Montage of landscapes and objects (e.g., houses and trucks) in a foggy and dusty atmosphere, with freaky background music. |
| 4 | HO | Home Alone | Home Alone | ha | 222 | 3:42 | A boy walks through his parents' house looking for his family, realising that he is home alone. |
| 4 | HO | Brockovich | Erin Brockovich | bv | 220 | 3:40 | A woman visits another woman asking her to testify at court. Afterwards she visits her place of work, a legal office, accompanied by her children. |
| 4 | HO | Star Wars | The Empire Strikes Back | sw | 246 | 4:06 | A man riding through an icy landscape gets attacked by a beast. People and imaginary animals work on aircrafts. A man and a woman are fighting. |

